# Nitric oxide S-nitrosylates CSF1R to augment the action of CSF1R inhibition against castration resistant prostate cancer

**DOI:** 10.1101/2022.06.09.495543

**Authors:** Manish Kuchakulla, Fakiha Firdaus, Rehana Qureshi, Yash Soni, Derek J Van Booven, Khushi Shah, Raul Ariel Dulce, Thomas Masterson, Omar Joel Rosete, Joshua M. Hare, Ranjith Ramasamy, Himanshu Arora

## Abstract

During progression of prostate cancer, sustained oxidative overload in cancer cells potentiates the overall tumor microenvironment (TME). Targeting the TME using colony-stimulating factor 1 receptor (CSF1R) inhibition is a promising therapy for castration-resistant prostate cancer (CRPC). However, the therapeutic response to sustained CSF1R blockade therapy (CSF1Ri) is limited as a monotherapy. We postulated that one of the causative agents for reduced efficacy of CSF1Ri and increased oxidation in CRPC is endothelial nitric oxide syntheses (eNOS). Results showed that in high grade PCa human specimens, eNOS is positively correlated with CSF1-CSF1R signaling and remains in an un-coupled state. The uncoupling disables eNOS to generate sufficient Nitric oxide (NO) that are required for inducing effective S-nitrosylation of CSF1R molecule at specific cysteine sites (Cys 224, Cys 278 and Cys 830). Importantly, we found that S-nitrosylation of CSF1R molecule at Cys 224, Cys 278 and Cys 830 sites is necessary for effective inhibition of tumor promoting cytokines (which are downstream of CSF1-CSF1R signaling) by CSF1R blockade. In this context, we studied if exogenous NO treatment could rescue the side effects of eNOS uncoupling. Results showed that exogenous NO treatment (using S-nitrosoglutathione (GSNO)) is effective in not only inducing S-Nitrosylation of CSF1R molecule, but it helps in rescuing the excess oxidation in tumor regions, reducing overall tumor burden, suppresses the tumor promoting cytokines which are ineffectively suppressed by CSF1R blockade. Together these results postulated that NO therapy could act as an effective combinatorial partner with CSF1R blockade against CRPC. In this context, results demonstrated that exogenous NO treatment successfully augment the anti-tumor ability of CSF1Ri in murine models of CRPC. Importantly, the overall tumor reduction was most effective in NO-CSF1Ri therapy compared to NO or CSF1Ri mono therapies. Moreover, Immunophenotyping of tumor grafts showed that the NO-CSF1Ri combination significantly decreased intratumoral percentage of anti-inflammatory macrophages, myeloid derived progenitor cells and increased the percentage of pro-inflammatory macrophages, cytotoxic T lymphocytes, and effector T cells respectively. Together, our study suggests that the NO-CSF1Ri combination has the potential to act as a therapeutic agent that restore control over TME and improve the outcomes of PCa patients.

## Introduction

Prostate cancer (PCa) is the most diagnosed non-skin malignancy in men^1^. Hormonal therapy is the treatment of choice for advanced PCa ^2, 3, 4, 5, 6^. However, due to moderate rate of success a significant number of patients progress to castration-resistant prostate cancer (CRPC) stage. This limited efficacy is believed to result in part from the unique ability of PCa to evolve through as-yet-unclear mechanisms in the tumor microenvironment (TME) that promote immune escape. TME is comprised of a variety of cell types of which tumor-associated macrophages (TAMs)^7^ frequently make up a substantial proportion and consists of two opposing phenotypes like classically activated (M1-like) and alternatively activated (M2 like), which have been correlated with anti and protumoral functions and with patient survival^8^. M1/M2 dichotomy is modulated via colony stimulating factor 1 (CSF1) which binds to CSF1 receptor (CSF1R) to control proliferation, differentiation, and survival of macrophages^9, 10^. Studies suggests that blocking CSF1R could delay tumor growth via TAM reduction^11, 12, 13, 14, 15^. However, single agent CSF1R blockade showed underwhelming results in phase 2 and 3 trials. One of the reasons for the limited efficacy is the feedback mechanism induced by activated CSF1R (in tumors) by recruitment of immunosuppressive and pro-tumoral TAMs and cytokines that are conducive to immune suppression^16^ and are not effectively targeted by single agent CSF1R blockade. Therefore, suggesting the need to finding ways to keep a check on TAMs and cytokines that derail the efficiency of CSF1R blockade therapy.

In this context, another relevant yet understudied aspect is Nitric oxide synthase (NOS). In mammals, three NOS isoenzymes are found. Of these, Neuronal and endothelial NOS (nNOS/NOS1 and eNOS/ NOS3, respectively) are constitutively expressed, produce Nitric Oxide (NO)(nM), and are regulated by Ca^2+^ binding to calmodulin ^17, 18, 19, 20^. NO has been shown to be involved in the regulation of adaptive immune responses by modulating T-cell activation, differentiation, promoting T-cell receptor mediated signaling from the immune synapse and M1/M2 macrophage polarization^17, 21^. The effect of NO are mechanistically imposed by covalent attachment of a nitroso group to a cysteine thiol (Protein S-nitrosylation) ^22^. NOSs of tumor cells, in contrast, synthesize superoxide and peroxynitrite which results in reduced tetrahydrobiopterin: dihydrobiopterin ratio (BH4:BH2). The reduced BH4:BH2 ratio results in uncoupling of NOSs and is observed in multiple cancer types^23^. One of the major impact of NOS uncoupling is the reduction in NO levels and increase in oxidative stress. Both of these aids the pro-tumorigenic cytokines such as NFkB^23^, IFNγ^24^, and TNFα^25^ which could further induce CSF1 expression in various cell types of TME and are not effectively targeted by single agent CSF1R blockade. Together, these findings warrant to study the impact of uncoupling of NOSs in conjunction with CSF1-CSF1R mediated M1/M2 dichotomy and TME in PCa which is unknown.

In this study, we found that eNOS remains in uncoupled state in CRPC. eNOS uncoupling results in reduced NO levels and increased oxidative stress which eventually results in keeping the cystine residues on CSF1R molecule in oxidized state. This in-turn negatively influences the anti-tumor effectiveness of CSF1R blockade. Therefore, we hypothesize that concomitant blockade of CSF1/CSF1R pathway in conjunction with exogenous increase in NO levels may improve immune function and CRPC treatment.

## Materials and Methods

### Human Samples

Patients enrolled in MD-SELECT/EDRN trial were used to collect whole blood samples. The patients were classified into different Gleason grades (6, 7 and 9) according to their PSA levels and MRI scans results. We took 3 samples corresponding to each Gleason grade. All human investigations were carried out after the IRB approval by a local Human Investigations Committee and in accord with an assurance filed with and approved by the Department of Health and Human Services. Data has been anonymized to protect the privacy of the participants. Investigators obtained informed consent from each participant. The whole blood was centrifuged at 3000 rpm to collect the white layer containing Peripheral blood mononuclear cells (PBMCs). The PBMCs were later lysed in RIPA buffer to check eNOS and CSF1 expression using ELISA. The flash frozen prostate biopsies (n = 5) corresponding to Gleason grade 6 and 9 were taken from the Cancer Modeling Shared Source (CMSR) at the University of Miami. The frozen biopsies were used for doing BH4 estimation and Griess test. The CMSR also provided with the biopsy sections which were used for doing immunohistochemical staining for eNOS, CD206, CSF1 and CSF1R.

### Cell culture

22R-v1 (CRL-2505), LNCaP <u>(CRL-1740)</u> and U937 cells (CRL-1593.2) were purchased from ATCC and maintained in RPMI 1640 medium supplemented with 10% fetal bovine serum, 100 units/ml penicillin, 100 mg/ml streptomycin and 2mM-L glutamine. TRAMPC2 cells (CRL-2731, purchased from ATCC) were maintained in DMEM medium supplemented with 5% fetal bovine serum, 100 units/ml penicillin and100 mg/ml streptomycin.

### Preparation of RNA and quantitative real-time PCR

Total RNA was extracted from cells using the TRIzol method and then reverse transcribed to complementary DNA using High-Capacity cDNA Reverse Transcription Kits (Applied Biosystems, USA) according to the manufacturer’s protocol. The quantitative RT-PCR for indicated genes was performed in SYBR Universal PCR Master Mix (BIORAD, USA). Quantitation of mRNAs was performed using BIORAD™ Gene Expression Assays according to the manufacturer’s protocol. Samples were analyzed using the BIORAD sequence detection system. All PCRs were performed in triplicate, and the specificity of the reaction was determined by melting curve analysis at the dissociation stage. The relative quantitative method was used for the quantitative analysis. The calibrator was the average ΔCt from the untreated cells. The endogenous control was glyceraldehyde 3-phosphate dehydrogenase (GAPDH).

### Western blotting

Cells were harvested and lysed in NP-40 buffer containing phenyl methyl sulfonyl fluoride and Protease Inhibitor Cocktail (Sigma, St. Louis, MO, USA). Protein expression was studied by exposing the membranes to antibodies against AR (Abcam, ab74272), AR-V7 (GeneTex, GTX33604), pERK (Cell signaling, 4370), ERK (Cell signaling, 9102S), pGSK-3beta (Cell signaling, 5558), p90RSK (Cell signaling, 11989), CD206 (Abcam, ab64693), VEGF (Abcam, ab46154) and GAPDH (Santa Cruz Biotechnology, SC47724). Immunoreactive bands were visualized using the Thermo Scientific Chemiluminescent Pico Kit.

### Immunohistochemistry and fluorescence staining

For immunohistochemistry, tissue sections were stained with hematoxylin and eosin and analyzed by a genitourinary pathologist. For fluorescence staining, tissue slides were processed using antibody against F4/80 (cell signaling, 70076), iNOS (abcam, 178945) and pERK (Cell signaling, 4370), followed by secondary antibodies tagged with Alexa Fluor® 488 or Alexa Fluor® 568 at room temperature for 30 minutes and 4,6-diamidino-2-phenylindole (Santa Cruz). All samples were assessed under a fluorescence microscope (Leica Microsystem, Wetzlar, Germany) at 60x magnification. Images were acquired using MetaMorph version 4.6 (Molecular Devices, Sunnyvale, CA, USA). Tumor xenograft tissues were fixed in 10% buffered formalin and embedded in paraffin. 5um thick sections were deparaffinized and rehydrated in sequential xylene and graded ethanol. Antigen retrieval was performed in 10 mM citrate buffer (pH 6.0) in a microwave oven. Peroxidase and non-specific protein blocking were done as per the instructions using the Abcam ABC detection kit (ab64264) and incubated with the following primary antibody dilutions: anti-Androgen Receptor (Abcam, ab74272), anti-Ki67 (Abcam, ab15580), anti-F4/80 (Cell signaling, 70076), anti-ARV7 (GeneTex, GTX33604) and anti-CD206 (Abcam, ab64693) with 1:150 dilution while a dilution of 1:50 was used for anti-Prostate-Specific Antigen (Santa-Cruz Biotech, sc7316). They were subsequently incubated with biotinylated goat anti-polyvalent secondary antibody, followed by development using DAB substrate as per the instructions on the kit. All sections were lightly counterstained with hematoxylin and mounted with Cytoseal XYL. Images were taken using a brightfield microscope (Nikon E200) at 10X and 40X magnification and quantification was done using ImageJ software (NIH, USA).

For fluorescence staining, post-antigen retrieval, the slides were permeabilized using 0.4% triton-X with 5%BSA. After permeabilization, sections were put in a blocking solution containing 5%BSA in PBST for 30 minutes at room temperature. Afterwards, incubation in primary antibodies were done overnight. Next day, slides were washed and incubated with standard fluorescent tagged secondary antibodies for an hour at room temperature. Following washing, antifade containing 4′,6-diamidino-2-phenylindole (DAPI) was used for mounting. Sections were imaged using a confocal microscope (Leica Microsystem, Wetzlar, Germany).

### MTT assay

22RV1 and LNCAP cells were seeded in 96-wells plate in quadruplets and were treated with varying doses of GSNO or GW2580 (CSF1R inhibitor) or with a combination of GSNO and GW2580. MTT (3-(4,5-dimethylthiazole)-2,5-diphenyltetrazolium bromide) assay reagents were added, and the absorbance was measured at 562nm after 0, 3, 5, 7, and 9 days.

### Animals

The animal protocol was approved by the Institutional Animal Care and Use Committee of University of Miami Miller School of Medicine, Miami, FL. SCID and C57BL6 (6 weeks old) mice were purchased from Jackson Laboratories (Bar Harbor, Maine). Castration experiments were performed in all the C57BL6 mice. For castration, mice were anesthetized using Isoflurane (Abbott Laboratories). The perineal region was cleaned three times with ethanol and a betadine scrub (VWR, AJ159778), and sterile dissecting shears were used to make a 4–5 mm incision in this region. Using two sterile forceps, the testes were located, and a ligature was made around the testicular vessels and the tunica albuginea that encases the testes. The testes were amputated with dissecting shears and the scrotum was sutured closed with 6-0 Ethicon black monofilament nylon (Ethicon Inc., 1665). A local triple antibiotic was applied over the region of the wound to facilitate healing. C57BL6 mice were grouped into control and experimental groups. All the mice were grafted with 1 million TRAMPC2 cells subcutaneously. The experimental group received 10mg/kg/day of GSNO, or 40mg/kg/day or 10mg/kg/day GSNO+40mg/kg/day GW2580 treatment intraperitoneally (IP) for 2 weeks, while the control group received PBS IP. After treatment, animals were survived for an additional two weeks before humanely sacrificing them. At the end time point, blood was collected via cardiac puncture and tumor grafts, lungs, and spleen were harvested for further analysis. Tumor volume (V) was measured regularly until the mice were sacrificed by measuring the length (L) and width (W) of the tumor with calipers by using the formula: V= 1/2(length × width^2^).

### Flow cytometry analysis for immune markers

For making single cell suspension from TRAMPC2 tumors, a piece of tumor was taken and minced with scissors or blade. These minced tumors were digested using digestion buffer containing collagenase by agitation at 37°C for 1.5 hrs and vortexing for a min, every 30 min. It was followed by adding RBC lysis buffer and 0.25%trypsin-EDTA and incubating again @ 37°C for 15 mins. Lastly, 5ml of chilled FACS buffer (PBS+2% FBS) was added and the digestion mixture was then filtered with 40um cell strainer. The cells were allowed to spin down at 450g for 5min and the collected pellet was resuspended in 2 ml FACS buffer. These isolated single cells were incubated with different antibodies following standard staining protocols for intra-cellular and membrane bound antibodies. The details of the antibodies are mentioned in Supplementary Table 2. Following staining, the cells were fixed in standard fixing solution containing 4% paraformaldehyde and taken for flow cytometry. The analysis was done using CYTEK Aurora flow cytometer.

### Griess Test

20 μL of Griess Reagent (Thermo Fischer Scientific G-7921) and 150 μL of the nitrite-containing sample were mixed in a microplate (sample capacity at least 300 μL per well). The mixture was incubated for 30 minutes at room temperature in dark. To prepare a photometric reference sample, 20 μL of Griess Reagent was mixed with 280 μL of deionized water. The absorbance of the nitrite-containing samples relative to the reference sample were measured in a spectrophotometric microplate reader at 548 nm. To convert absorbance readings to nitrite concentrations, the method recommended by the manufacturer was used.

### Tetrahydrobiopterin (BH4) ELISA

The kit (Catalog No. ABIN6957559, Antibodies online) is a competitive inhibition enzyme immunoassay technique for the in vitro quantitative measurement of tetrahydrobiopterin in serum, plasma, tissue homogenates, cell lysates, cell culture supernatants. The amount of lysates used was 100ug of protein.

### Human eNOS/NOS3 ELISA

Peripheral blood mononuclear cells (PBMCs) collected from patient blood samples were used for the estimation of NOS3 levels (Catalog # EH169RB, Thermo Fischer Scientific, USA). The steps followed were according to the kit instructions. 100ug of cell lysate was used for quantification.

### M-CSF (CSF-1) Human ELISA

Peripheral blood mononuclear cells (PBMCs) collected from patient blood samples were used for the estimation of CSF-1 levels (Catalog # EHCSF1, Thermo Fischer Scientific, USA). 100ug of cell lysate was used for the assay. The steps followed are the same as mentioned in the instruction manual.

### Cytokine antibody array

Proteins were isolated from tumors of the CRPC mice that had received treatment with PBS and GSNO. After quantification using Bradford assay, 2.5mg/ml protein lysates were screened for secreted protein using RayBio Human Cytokine Array C5, Code: AAH-CYT-5-2 (RayBiotech, Norcross, GA, USA) according to the manufacturer’s instructions. The blots were analyzed using ImageJ software (National Institutes of Health, Bethesda, MD, USA). A total of 80 molecules were selected for detection namely: ENA-78, G-CSF, GM-CSF, GRO, GRO-alpha, I-309, IL-1alpha, IL-1beta, IL-2, IL-3, IL-4, IL-5, IL-6, IL-7, IL-8,IL-10, IL12-p40, IL-13, IL-15, IFN-gamma, MCP-1, MCP-2, MCP-3, M-CSF, MDC, MIG, MIP-1 beta, MIP-1-delta, RANTES, SCF, SDF-1, TARC, TGF-beta 1, TNF-alpha, TNF-beta, EGF, IGF-1, Angiogenin, Oncostatin M, TPO, VEGF, PDGF-BB, Leptin, BDNF, BLC, CK beta 8-1, Eotaxin, Eotaxin-2, Eotaxin-3, FGF-4, FGF-6, FGF-7, FGF-9, Flt-3 Ligand, Fractalkine, GCP-2, GDNF, HGF, IGFBP-1, IGFBP-2, IGFBP-3, IGFBP-4, IL-16, IP-10, LIF, LIGHT, MCP-4, MIF, MIP-3-alpha, NAP-2, NT-3, NT-4, Osteopontin, Osteoprotegerin, PARC, PIGF, TGF- b 2, TGF- b 3, TIMP-1, and TIMP-2 respectively.

### Biotin switch assay and SNO protein purification

The S-nitrosylated proteins were visualized by biotin-switch assay following the manufacturer’s guidelines (S-Nitrosylated Protein Detection Kit (Biotin Switch), Item No. 10006518, Cayman Chemical, Ann Arbor, MI, USA). A small piece of tumor tissue or cell pellet was taken and washed twice with Wash Buffer. The pellets were resuspended in “Buffer A containing Blocking Reagent” and incubated for 30 min at 4 °C with shaking. The incubated samples were centrifuged at 130,000×rpm for 10 min at 4 °C, and the supernatant was transferred to 15 ml centrifuge tubes. Two milliliters of ice-cold acetone was added to each sample, and the mixture was incubated at − 20 °C for at least 1 h. The protein of each sample was pelleted by centrifugation for 10 min at 4 °C. “Buffer B containing Reducing and Labeling Reagents” was added to resuspend the proteins, with incubation for 1 h at room temperature. The biotinylated protein was precipitated by acetone and rehydrated with the appropriate amount of Wash Buffer.

A total of 40ug of protein was used for labeling and running the standard western blot for detecting the nitrosylated protein.

### RNA Sequencing and Enrichment analysis

Fastq files were downloaded from Illumina’s BaseSpace cloud application. FastQC was performed on the fastq files to ensure acceptable base quality and GC content and to check for adapters. Adapters were trimmed using the Trim_Galore software to remove the Illumina Universal Adapter. After adapter trimming, Fastq_screen was performed to check for bacterial contamination. Once no contamination was confirmed, general alignment was performed using the STAR RNAseq aligner to the hg19 genome. During this alignment, raw counts were produced against the GENCODE gene features (v19) and reformatted into a matrix for further statistical processing. After alignment, PicardTools was used to calculate sample by sample alignment quality metrics statistics. Differential expression was performed on the raw count matrix using the edgeR software. Statistical significance was defined as any gene feature that had an FDR value of below 0.05. Gene set enrichment analysis was performed by means of the Broad’s GSEA executable jar file using the MSigDB as the reference database. Normalized log2CPM values were used as the expression values. These CPM values were filtered by log2CPM of 1 to limit the noise of the data that was used for enrichment. Three sets of runs were completed against hallmark genes (h).

### Activation of Monocytes for M1/M2 polarization

U937 cells were activated using different cytokines as done by previous studies. The cells were treated in RPMI 1640 medium supplemented with 5% cFBS, in 100mm flat-bottom culture plates. Activation treatments consisted of (1) no stimulation control (mock); (2) PMA 20 ng/mL for 48 h (PMA-only control); (3) pre-treatment with PMA 20 ng/mL for 48 h, followed by LPS 50 ng/mL and IFN-γ10 ng/mL for 48 h (condition favoring M1 polarization, M1 cocktail); (4) pre-treatment with PMA 20 ng/mL for 48 h, followed by IL-4 25 ng/mL and IL-13 25 ng/mL for 48 h (condition favoring M2 polarization, M2 cocktail). In addition to these treatments, three drug combinations, i.e., GSNO (50uM), CSF1Ri (0.5uM) and a combination of both GSNO and CSF1Ri were also used to check their effect on monocyte activation as well as macrophage polarization.

### Macrophage polarization study for studying the interaction of AR and GSNO

For this study, U937 cells (purchased from ATCC) were used as a model for monocyte-macrophage differentiation. The study was divided into two steps: a) Collection of conditioned media and cell lysates from 22Rv1 cells and b) Treatment of U937 cells with conditioned media from 22Rv1. For preparing the 22Rv1 cell conditioned media (CM), 3.0 × 105 cells were seeded into 6-well culture plates and allowed to adhere overnight. Next day, the cells were starved using charcoal stripped FBS for about 12 hours and then 2 ml of medium containing 1% FBS. The cells were treated with 4μM enzalutamide (ENZA, Selleckchem, MDV3100) in the presence or absence of 50 μM GSNO. The untreated 22Rv1 cells served as control. The media was collected 48 h later. The supernatant was centrifuged at 1800 rpm for 10 min and was collected as CM. U937 cells were treated with the collected CM from these different treatment conditions. The cells were collected at 48 hrs for analysis of M2 macrophage marker, CD206 (Abcam, ab64693) and AR (Abcam, ab74272).

### Organoid cultures using 22Rv1 cells for Immunohistochemical (IHC) staining

The organoid cultures using 22Rv1 cells were generated using the protocol followed by Ma et al (2017). A total of 250,000 cells/ml were suspended in thawed Matrigel and a drop was added in each well of a 24 well plate. Organoids were maintained in adDMEM/F12 media with growth factors and 10 μM Y-27632 dihydrochloride. The media was refreshed every 2-3 days. From 7 days after initial plating, an organoid culture medium without Y-27632 dihydrochloride was used. The culture was ended on day 14. Later the organoids were fixed in 4% paraformaldehyde and processed for paraffin embedding and sectioning.

### Site directed mutagenesis

CSF1R ORF clone (OHu24034, NM_001349736.1) was procured from GenScript Biotech (NJ, USA). Site directed mutagenic changes were done to incorporate 3 cysteine deletions at C224, C278 and C830 (as determined by GPS-SNO 1.0 software with high threshold) using Quick Change Lightning Multi Site-Directed Mutagenesis Kit (Agilent Technologies, USA). Briefly, 40 ng plasmid was subjected to PCR amplification as per standard kit guidelines using mutagenic primers 5’-tgcccagatcgtgtcagccagcagcg-3’, 5’-cgctgctggctgacacgatctgggca-3’; 5’-cgttgctggccacggagtagttgccg-3’, 5’-cggcaactactccgtggccagcaacg-3’ and 5’-ctctgaaccgtgtagacgtcaaagatgctctctg-3’, 5’-cagagagcatctttgacgtctacacggttcagag-3’ for cysteine del1, del2 and del3 respectively. Following PCR, 10ul of the product was subject to DpnI digestion for 5min at 37oC and transformed into chemically competent DH5α cells (NEB, USA) by heat shock at 42oC for 30 sec. Resulting transformants were grown in SOC media for 1hr at 37oC and selected overnight on LB agar plates containing 100μg/ml Ampicillin. Following day, single colonies were selected and further grown in LB broth containing 100μg/ml Ampicillin for 12 hours. Plasmid isolation and purification was done using plasmid miniprep kit (Qiagen, Germany) as per standard instructions. Sanger sequencing for confirmation of deletion was done by Genewiz, USA. For verification of these deletions, the confirmed wild type as well as mutant clones were transfected in 22Rv1 cells using Lipofectamine 3000 reagent. After transfection, cells were treated with/without 50μM GSNO and cells were collected after 48 hrs to do western blots using anti-Androgen Receptor (Abcam, ab74272) and anti-beta Actin antibody (Cell Signaling Technology, 4970) antibodies.

### Statistical analysis and calculation of sample size

GraphPad Prism (GraphPad Software) was used for statistical analysis. All data are presented as means ± SEM. The statistical significance between two groups was determined by unpaired two-tailed *t* test. Multiple group comparisons were performed using a one-way analysis of variance with Tukey least significant difference test. In all cases, *p* < 0.05 was considered statistically significant.

## Results

### Prostate Cancer progression is associated with increased CSF1 concentration and Nitric oxide synthases

The expression of CSF1 and NOSs (eNOS, iNOS and nNOS) was studied using RNA sequencing data for Prostate adenocarcinoma from The Cancer Genome Atlas (TCGA). Results showed that, with increase in cancer grade (Gleason 6-10), the expression of CSF1 and eNOS increases (Figure 1A, Supplementary Figure 1A). Additionally, the Peripheral blood mononuclear cells (PBMCs) isolated from the blood of healthy individuals and patients with primary (Gleason 6), mid (Gleeson 7) or distant metastasis (Gleason 9) PCa (n=2 per condition), staged according to American committee on Cancer was used to evaluate the expression of CSF1 and eNOS. Both CSF1 and eNOS were significantly increased (p<0.05) in high grade PCa patients (Figure 1B-C, Supplementary Figure 1B). We also evaluated the expression of eNOS in patient biopsies from Gleason 6 and Gleason 9 (n=3); and found an overall increase in its expression (***p ≤0.001) (Supplementary Figure 1C). These findings indicate that CSF1 and eNOS production is elevated with disease progression.

**Fig 1:**
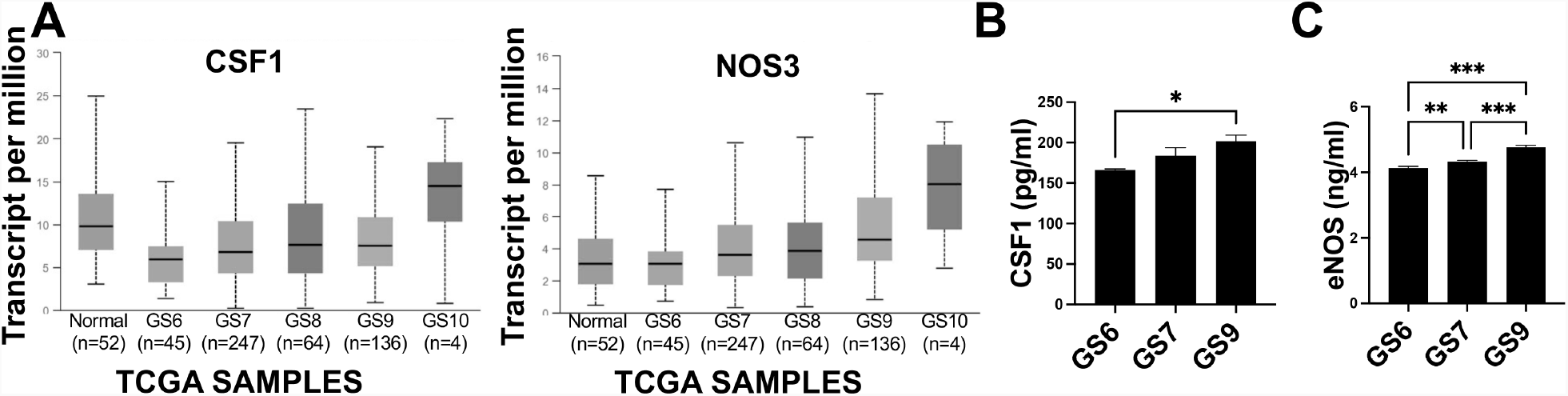
Prostate Cancer progression is associated with increased CSF1 concentration and Nitric oxide synthases. The relative expression of CSF1 and NOS3 was studied using RNA sequencing data for Prostate adenocarcinoma from The Cancer Genome Atlas (TCGA). (A) Graphs showing expression of CSF1 and NOS3 across different Gleason Grades. (B) Quantification of CSF1 and eNOS was done by enzyme-linked immunosorbent assay (ELISA) performed in PBMCs collected from different patient blood samples with different Gleason grades (n=2). CSF1 levels are expressed as pg/ml while eNOS levels are expressed as ng/ml. Data are means ± SEM. n, number of samples. GS, Gleason grade. *p < 0.05; **p < 0.01; ***p < 0.001.

### High number of eNOS^+^ cells in high grade PCa are correlated with enrichment of CSF1^+^, CSF1R^+^ cells

High expression of CSF1 and CSF1R were found to be specific to cancer regions in high-grade PCa (e.g., Gleason 9) in comparison to normal adjacent regions (n=3 patients) (Supplementary Figure 2A). Additionally, with increasing grade of PCa (Gleason 6, 7, and 9), expression of anti-inflammatory (M2) macrophages (CD206) increase (Supplementary Figure 2B). To determine whether the expression of eNOS is correlated with CSF1, CSF1R and CD206, chromogenic immunohistochemistry (IHC) was performed using specimens from normal adjacent and metastatic regions of high grade PCa (n=3). Results showed that high grade cancer regions with higher expression of eNOS had high expression of CSF1, CSF1R and CD206 (Figure 2A). Furthermore, RNA sequencing analysis from TCGA data showed, the expression of eNOS/NOS3 is positively correlated with the expression of CSF1, CSF1R, CD163 and several other tumors promoting immune cell markers in high grade PCa (Gleason 9)(Figure 2B). Moreover, immunofluorescence staining (IF) demonstrated that positive correlation between eNOS and CSF1 is specific to high grade PCa (n=3 specimen for Gleason grade 6 and 9)(Figure 2C). Together these results strongly argue that the abundance of eNOS positively correlates with TAM in human PCa.

**Fig 2:**
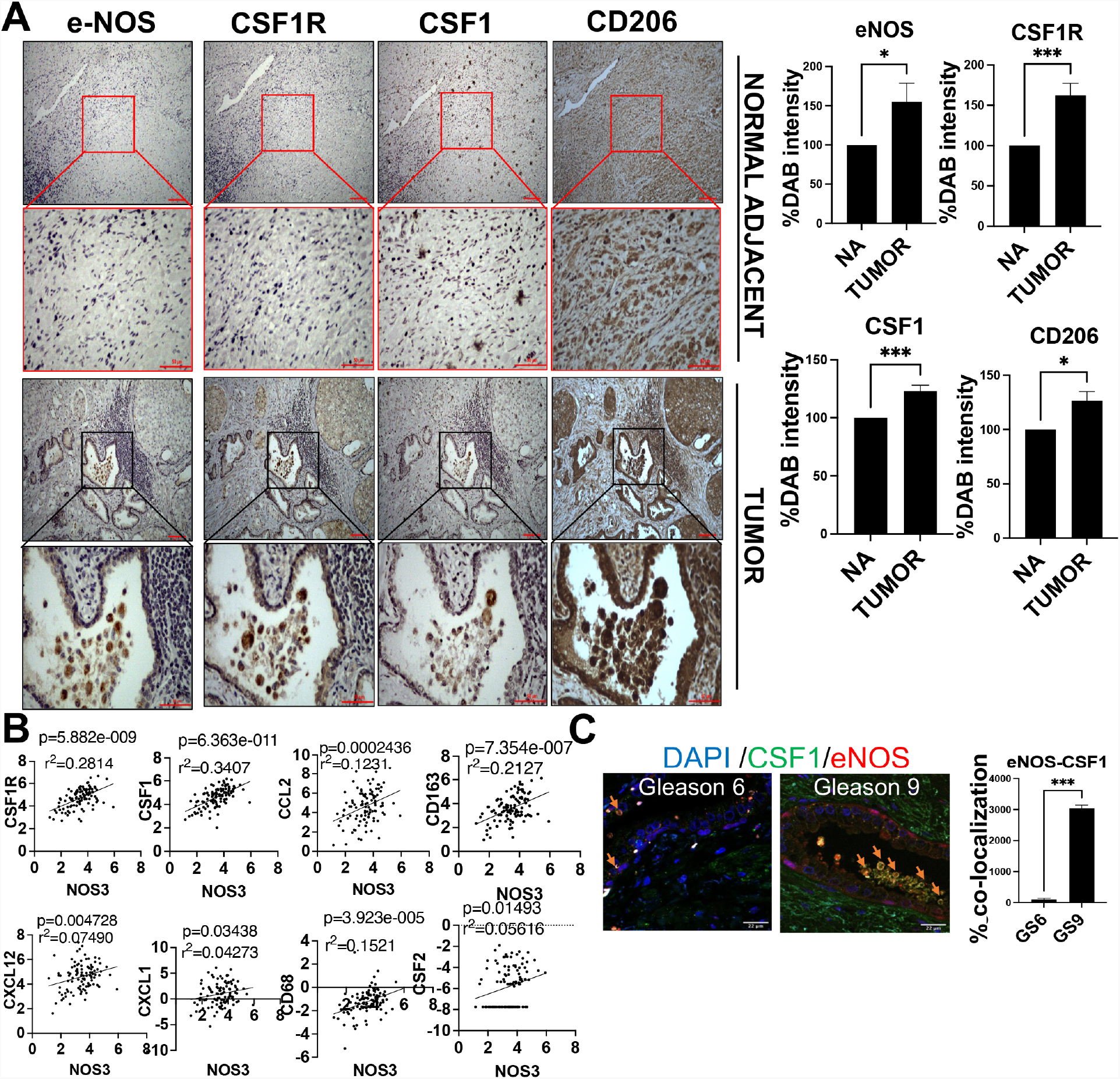
eNOS expression in prostate cancer biopsies is correlated with enrichment of CSF1^+^, CSF1R^+^, and CD206^+^ cells. (A) Representative images of eNOS, CSF1R, CSF1, and CD206 immunostaining selected from a tumor region with either high or low eNOS expression in a patient biopsy sample. The red and blue square in the upper panel indicates a region of interest that has been magnified in the lower panels. Scale bar, 100 μm. (B) Matrix of scatterplots showing correlations between CD8A, CD8B, CSF1, CSF1R, CD68, and CD163 gene expression specific to high grade prostate cancer (Gleason grade 9) of TCGA. Correlation was assessed using Spearman’s correlation coefficient. Black lines indicate the local regression (LOESS) fit. P, P value; n, number of samples; r^2^, Spearman’s correlation coefficient. (C) Representative multiplexed fluorescence staining images of tumor tissue from one prostate biopsy patient stained with 4′,6-diamidino-2-phenylindole (DAPI) (blue) and antibodies against CSF1 (green) or eNOS (red). Percentage of co-localization for CSF1+eNOS+ cells are represented for the patients listed in table S1 (n = 3). Data are means ± SEM of three images per tumor area for each patient. *p < 0.05; ***p < 0.001.

### Tumor cells are a source of eNOS in human Prostate cancer

We then asked whether PCa cells could be a source of eNOS. We performed IF on histological sections of three Gleason 9 specimens. PCa cells were identified by their expression of the FOXM1^26^. We found that out of 73.77% FOXM1+ PCa cells, 54.28% expressed eNOS in all three patient biopsy sections (Figure 3A). We then assessed CSF1 in relation to FOXM1 cells abundance and found that the CSF1 expression was higher in regions with higher FOXM1 expression (Figure 3B).

**Fig 3:**
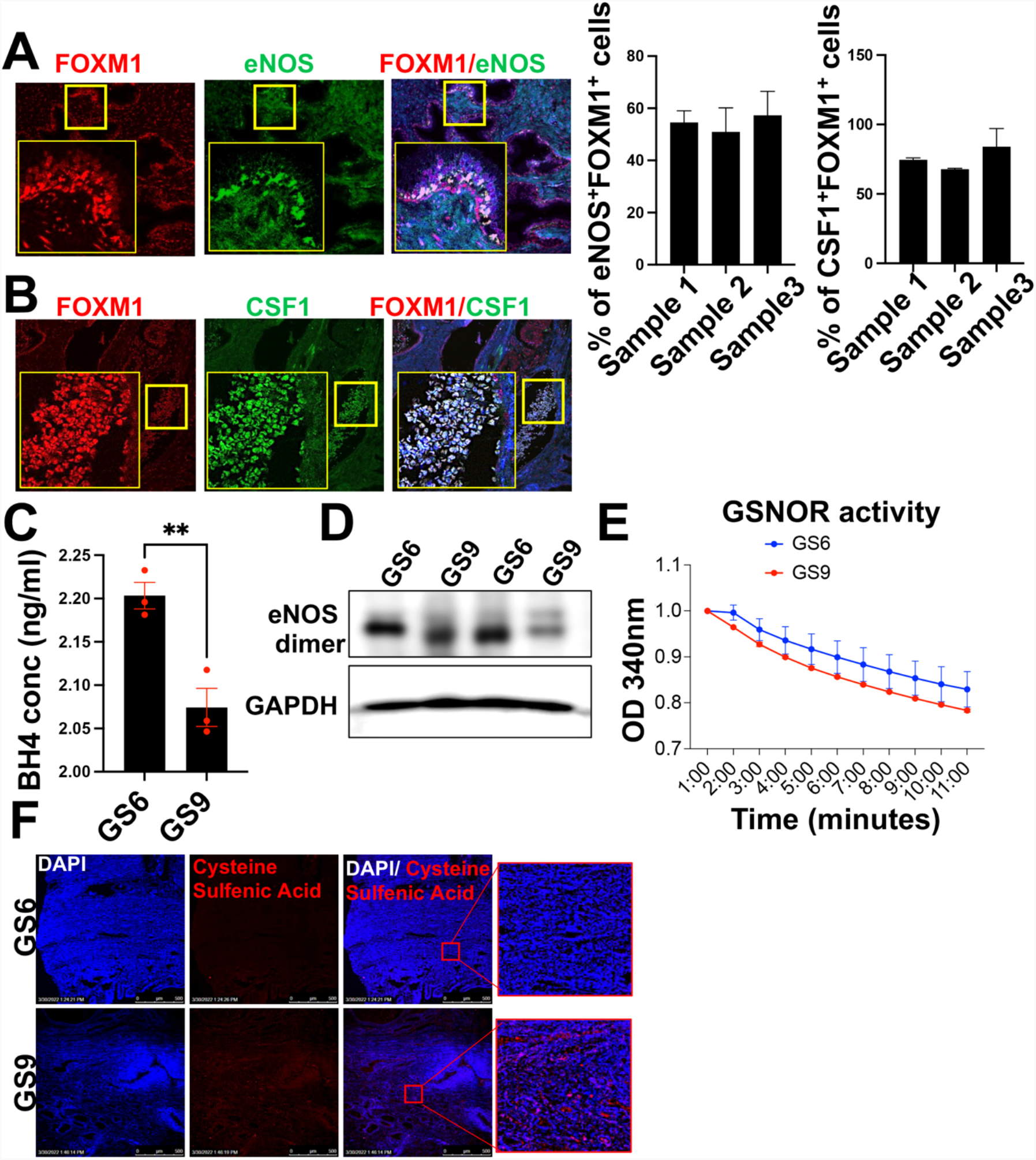
Tumor cells are a source of eNOS in human prostate cancer. (A-B) Representative multiplexed fluorescence staining images of tumor tissue from one prostate biopsy patient stained with 4′,6-diamidino-2-phenylindole (DAPI) (blue) and antibodies against FOXM1 (red) and eNOS/CSF1 (green). Percentage of FOXM1+ prostate cancer cells that are also eNOS+ as well as CSF1+ in prostate adenocarcinomas (n = 3) from the patients listed in table S1. Data are means ± SEM of three images per tumor area for each patient. Scale bar, 22 μm. (C) BH4 concentration estimated using competitive enzyme-linked immunosorbent assay (ELISA) using prostate biopsy lysates (n=3). **p < 0.01. Scale bar, 22 μm. (D) Uncoupling of eNOS as shown by dimers using prostate biopsies from different Gleason grades. The lysates were run at low temperature without beta-mercaptoethanol to look for dimers. (E) GSNOR activity measured and recorded at 340 nm for a span of 10 minutes in prostate biopsies from Gleason grade 6 and Gleason grade 9 (n=3). (F) Representative immunofluorescent images for patient biopsies (Gleason grade 6 and 9) stained with 4′,6-diamidino-2-phenylindole (DAPI) (blue) and Cysteine sulfinic acid (red). Scale bar 100um.

### eNOS in human Prostate cancer are in uncoupled state

The main and most important function of NOSs is synthetizing nitric oxide (NO). Reduced ratio BH4:BH2 in several cancer types, destabilizes the NOS subunits (uncoupling) switching the enzymatic activity toward a superoxide (O2^-^) generating enzyme^23, 27^. We checked BH4 to evaluate if, the eNOS in high grade human PCa are in coupled or uncoupled state. Results showed a significant reduction of BH4 in patients with distant metastasis compared to specimens from primary PCa (Gleason 6)(n=3 each)(Figure 3C). This was further reflected by the reduced dimerization of eNOS in primary PCa specimens compared to distant metastasis (Figure 3D). Next, we tested the activity of the S-Nitrosoglutathione reductase in proteins from primary and metastatic human PCa tissues (n=3 each). Results showed an enhanced GSNOR activity in metastatic specimens, which suggest a higher depletion rate of GSNO (Figure 3E). Therefore, reflecting that in high grade human PCa, eNOS becomes upregulated but uncoupled. These results, together with the increased GSNOR activity in these samples, suggest a profound depletion in intracellular NO availability.

### Uncoupled eNOS could negatively influence the anti-tumor effectiveness of CSF1R blockade therapy against CRPC

In the eNOS enzyme, NADPH-derived electrons flow from the reductase domain toward the oxygenase domain, and in the presence of BH4, these electrons react with oxygen and L-arginine and lead to the formation of L-citrulline and NO^28^. However, in the absence or reduced levels of BH4 (as observed in high grade PCa), oxidation of oxygen occurs, and the enzyme subsequently releases superoxide^29, 30^. In this context, considering a strong and positive correlation between eNOS and CSF1-CSF1R axis and based on findings which suggest that neo-adjuvant CSF1R inhibition is minimally effective against high grade PCa^31^, we hypothesize that increased oxidation because of uncoupling of eNOS could reduce the efficiency of CSF1R blockade. To study this, first we evaluated the extent of oxidation in histological sections from primary or metastatic PCa specimens (n=3 each) through IF with antibody against Cysteine Sulfenic Acid (CSA) ^32, 33^. Results showed an increase in oxidation pockets in high grade PCa (Figure 3F). Importantly, CSF1R expression was found to be localized in these oxidized pockets. Furthermore, we generated prostate organoids using 22RV1 (CRPC) cells. These organoids were exposed to single agent CSF1R blockade (GW2580). Immunostaining on organoid sections showed minimal changes in the expression of CSA or eNOS upon CSF1R blockade (Figure 4A). Next, we tested the effects of CSF1R blockade in CRPC in-vivo models (castrated C57BL6 mice were grafted with TRAMPC2 cells)(Figure 4B). Exposure to CSF1Ri (40mg/kg/day IP) was able to reduce the tumor burden (Figure 4C), but GSNOR activity remained unchanged (Figure 4D). Furthermore, like organoids, IHC on murine tumor grafts demonstrated no change in oxidation pockets in control or treatment group (Figure 4E). Together the results suggested that eNOSs is up-stream to CSF1-CSF1R axis and CSF1R blockade has minimal influence on oxidation. Furthermore, despite reduction in tumor burden upon CSF1R blockade, there was a minimal change in the expression of AR, pERK, p-GSK, VEGF, ARV7 and Ki67 (Supplementary Figure 3A-B). Moreover, cytokine array showed a minimal impact of CSF1R blockade on several tumor promoting cytokines such as ENA-78, BLC, IP-10, Oncostatin M, FGF-6, IGFBP-3, RANTES, Osteopontin, GCP-2, IL-15, MCP-4, NT-4, TGF-b 3, IL-3, SCF, IL12-p40, IGFBP-4, FGF-4, SDF-1, PDGF-BB, IL-1beta, PARC, TPO, IFN-gamma, FGF-7, and IL-2 respectively (Supplementary Figure 3C). The role of each of these cytokines is summarized in Supplementary Table 1.

**Fig 4:**
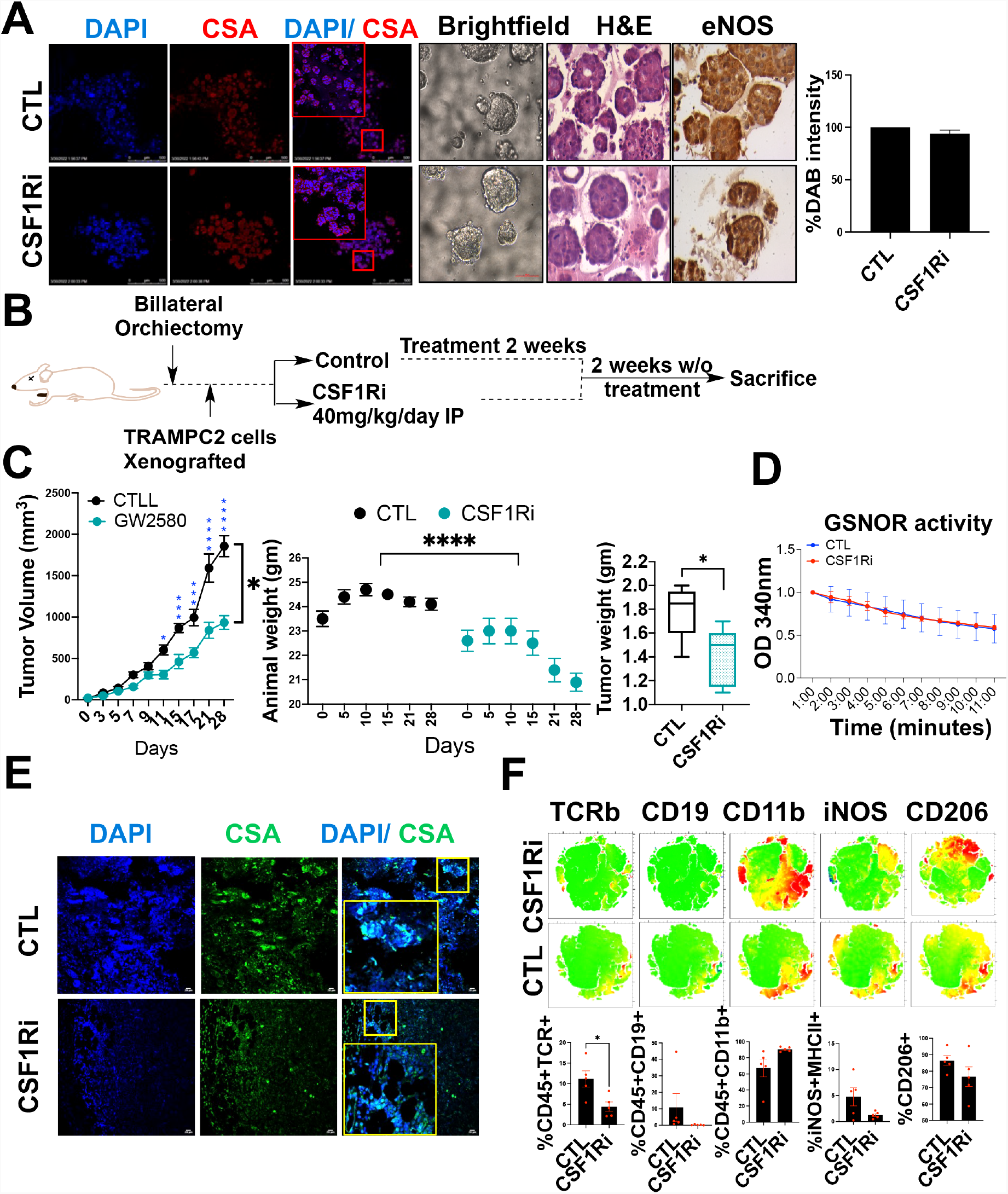
CSF1R inhibition reduces PCa tumor burden but potentiates pro-tumorigenic signatures. (A) Representative bright-field, H and E and immunofluorescent images for 22Rv1 derived organoids stained with 4′,6-diamidino-2-phenylindole (DAPI) (blue) and Cysteine sulfinic acid (red) showing oxidation status. The chromogenic substrate DAB (brown) represents the expression of eNOS in untreated control as well as CSF1Ri treated organoids. Scale bar 400um. (B) Experimental plan. (C) Mean change of tumor volume (±SEM; versus tumor volume at treatment start) in C5BL6 mice treated as indicated. Vehicle (n = 5), α-CSF1R (n = 5). Animal weight change across the course of experiment from Day 1 till Day 28. Tumor weight plotted as Mean±SEM (n=5) measured at the end of experiment. (D) GSNOR activity measured at 340nm for a span of 10 minutes using tumor lysates treated with vehicle and CSF1R inhibitor (n=3). (E) Representative immunofluorescent images for tumor sections for mice treated with vehicle and CSF1R inhibitor and stained with 4′,6-diamidino-2-phenylindole (DAPI) (blue) and Cysteine sulfinic acid (red) (n=3). (F) Representative tSNE plots and graphs for various immune cell surface markers (TCRβ, CD19, CD11b, iNOS and CD206) for tumor cells isolated from mice tumors treated with vehicle and CSF1R inhibitor

Next, tumor grafts from control and treatment groups were subjected to comprehensive immunophenotyping, which utilizes a standardized panel of antibodies against each of immune cell types (Supplementary Table 2). Results demonstrated minimal effects of CSF1R blockade on several of the cell types (CD206^+^ (%CD45^+^)(M2 macrophages), iNOS^+^/MHC-II^+^ cells (%CD45^+^)(M1 macrophages), CD11b^+^(%CD45^+^)(myeloid cells), % B cells and % T cells)(Figure 4F, Supplementary Figure 3D). Together the results suggested that the low efficiency of CSF1R blockade could potentially be accounted to increased oxidation and reduced NO levels which results from eNOS being in an un-coupled state.

### Exogenous induction of NO overcomes the negative effects of eNOS uncoupling and demonstrates tumor inhibitory role against CRPC

We asked if an exogenous induction of NO could rescue the negative effects of increased oxidation and eNOS uncoupling on CSF-CSF1R axis. For this we focused on CSF-CSF1R induced functions which primarily includes macrophage polarization, to promote angiogenesis, tissue remodeling, and immune suppression^34^. In this context, prostate organoids (from 22RV1 cells) were exposed to exogenous source of NO (S-Nitrosoglutathione (GSNO)). Results showed a significant reduction (**p≤0.01) in CD206 expression upon treatment (Figure 5A-B). Second, we used U937 cells (model for monocyte-macrophage differentiation) and treated them with Phorbol-12-Myristate-13-Acetate (PMA) in the presence of M1 or M2 macrophage inducing cocktails (Supplementary Figure 4A). These cells were then exposed to GSNO (50uM) and the percentage shift in the population of M1 (CD68^+^) or M2 (CD206^+^CD163^+^) cells was evaluated. Results showed a significant increase in M1 and a reduction in M2 macrophage populations (Supplementary Figure 4A), suggesting that exogenous increase in NO could inhibit macrophage polarization in-vitro. Furthermore, studies suggest that AR signaling in macrophages promotes PCa cell migration and invasion^35, 36, 37^. To study if the exogenous increase in NO could inhibit AR signaling in macrophages to influence macrophage polarization, we exposed U937 cells to conditioned media (CM) from 22RV1 cells which were treated with enzalutamide (AR antagonist) or with GSNO (Supplementary Figure 4B). Both, enzalutamide and GSNO treatment reduced the expression of AR and CD206 in 22RV1 and U937 cells (Supplementary Figure 4C). Third, we asked if exogenous induction of NO could inhibit tumor growth. For this, we tested the effects of GSNO on murine CRPC in-vivo models. Results showed that GSNO inhibited tumor burden by showing a significant decrease (*p≤0.05) in the tumor volume and tumor weight (Figure 5C). The tumor grafts from GSNO treatment group showed reduced oxidation pockets compared to control group (Figure 5D). Moreover, the expression of AR, ARV7, and Ki67 was reduced upon GSNO treatment (Supplementary Figure 5A). Furthermore, cytokine array showed that GSNO reduced the expression of ENA-78, BLC, IP-10, Oncostatin M, FGF-6, IGFBP-3, RANTES, Osteopontin, GCP-2, IL-15, MCP-4, NT-4, TGF-b 3, IL-3, SCF, IL12-p40, IGFBP-4, FGF-4, SDF-1, PDGF-BB, IL-1beta, PARC, TPO, IFN-gamma, FGF-7, and IL-2 (Supplementary Figure 5B). Next, we subjected the tumor grafts to immunophenotyping. Results showed that GSNO increased the percentage of iNOS^+^/MHC-II^+^ cells (%CD45^+^)(M1 macrophages), tumor infiltrating lymphocytes (CD8^+^(%CD45^+^) cells), CD44^+^CD62L^-^ cells (%CD8^+^)(EM CD8^+^), CD44^+^CD62L^+^ cells (%CD8^+^)(MEM CD8^+^), and naïve CD8^+^ cells (%CD45^+^) and reduced the percentage of CD206^+^ (%CD45^+^)(M2 macrophages), F480^+^ (%CD45^+^)(macrophages), and CD11b^+^(%CD45^+^)(myeloid) cells (Figure 5E). Together the results suggested that exogenous increase in NO could rescue increased oxidation; inhibit macrophage polarization; and have tumor inhibitory effects against CRPC.

**Fig 5:**
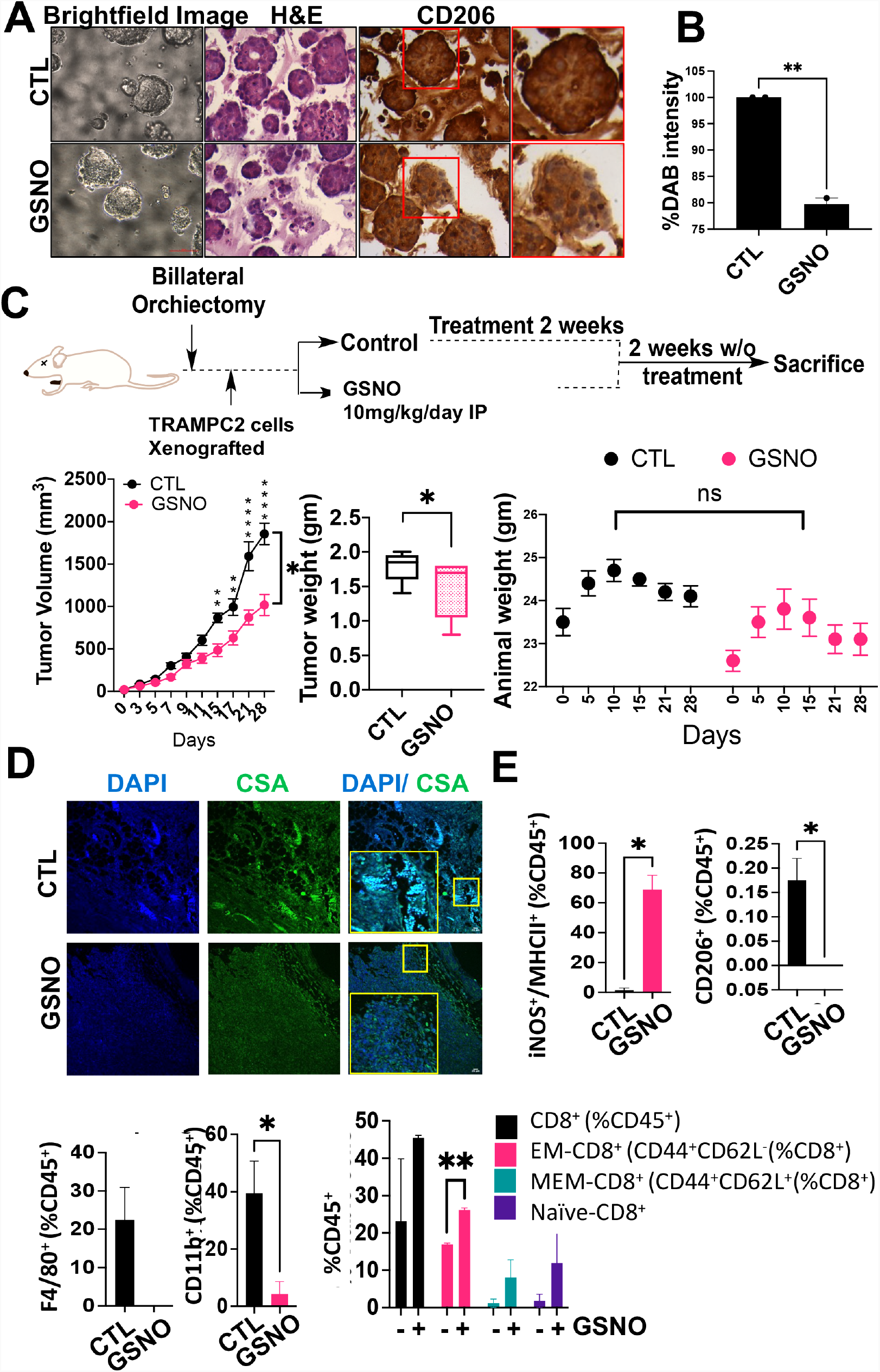
Increased NO reduces PCa tumor burden and pro-tumorigenic signatures. (A) Representative bright-field, H and E and immunofluorescent images and DAB chromogenic substrate showing expression of CD206 in 22Rv1 derived organoids. Scale bar 400um. (B) Graph showing %DAB intensity for CD206 expression. Data are ±SEM. Scale bar 400um. (C) Mean change of tumor volume (±SEM; versus tumor volume at treatment start) in C5BL6 mice treated as indicated in the flow diagram. Vehicle (n = 5), S-nitrosoglutathione, GSNO (n = 5). Animal weight plots across the course of experiment from Day 1 till Day 28. Tumor weight plotted as Mean±SEM (n=5) measured at the end of experiment. (D) Representative immunofluorescent images for tumor sections for mice treated with vehicle and GSNO and stained with 4′,6-diamidino-2-phenylindole (DAPI) (blue) and Cysteine sulfinic acid (red) (n=3). (E) Graphs showing the percentage of various immune cell populations in tumor grafts representing percentage of iNOS+/MHCII+ (M1-macrophages), CD206+, F4/80+ (M2-macrophages) and CD11b+ myeloid cells. Also, the cytotoxic T cell population (CD8+), EM-CD8+ (CD44+CD62L-), MEM-CD8+ (CD44+ CD62L+) and naïve CD8+ are represented in GSNO tumors treated as indicated and analyzed by flow cytometry (Mean±SEM). Statistical analysis by Student’s t test. ***P < 0.001; **P < 0.01; *P < 0.05. ns=not significant.

### Nitric oxide S-nitrosylates CSF1R to induce tumor inhibitory functions

One of the key mechanisms of action of GSNO is through the coupling of a nitroso moiety from NO-derived metabolites to a reactive cysteine leading to the formation of a S-nitrosothiol (SNO), a process commonly known as S-nitrosylation ^38^. Considering a strong correlation between un-coupled eNOS and CSF1-CSF1R signaling and inhibitory impact of both GSNO and CSF1R blockade against M1-M2 dichotomy, we hypothesized that GSNO could potentially S-nitrosylate CSF1R to inhibit CRPC tumor burden, overall oxidation and macrophage dichotomy. For this, biotin-switch assay was performed to measure S-Nitrosylated (S-NO) CSF1R proteins from tumors grafts from mice which were exposed to GSNO treatment. Results confirmed that GSNO treatment induced S-nitrosylation of heavy chain of CSF1R (Figure 6A). Next, we screened CSF1R to identify potential cystine sites using GPS-SNO 1.0 software which conformed to an acid-base nitrosylation conservative motif (14, 29). A total of 20 cystine sites were identified on CSF1R structure, with three sites Cys224, Cys278 and Cys830 having high predicted threshold (cutoff 2.443) for being S-nitrosylated (Figure 6B and Supplementary Table 3). Cysteine mutations were generated for Cys224, Cys278 and Cys830 by site-directed mutagenesis. A topology map of CSF1R indicated that the location of the 3 mutated cysteines in CSF1R structure is shown in Figure 6C. Next, we characterized the cellular localization of these mutants (Cys224, Cys278 and Cys830) in 22RV1 cells to determine whether cysteine mutations lead to a disruption of protein 3D structure. All the mutants localized primarily to the nucleus of the cells in a pattern like that of wild-type CSF1R protein (Supplementary Figure 6A). Further, we transfected the 22RV1 cells with the CSF1R plasmids (wild type and three mutants), followed by exposing the cells to vehicle or GSNO for 48 hours. Biotin-switch assay confirmed that reduced GSNOR activity was directly correlated with blocked Cys224, Cys278 and Cys830 sites which further inhibited the GSNO induced S-nitrosylation of CSF1R (Figure 6D).

**Fig 6:**
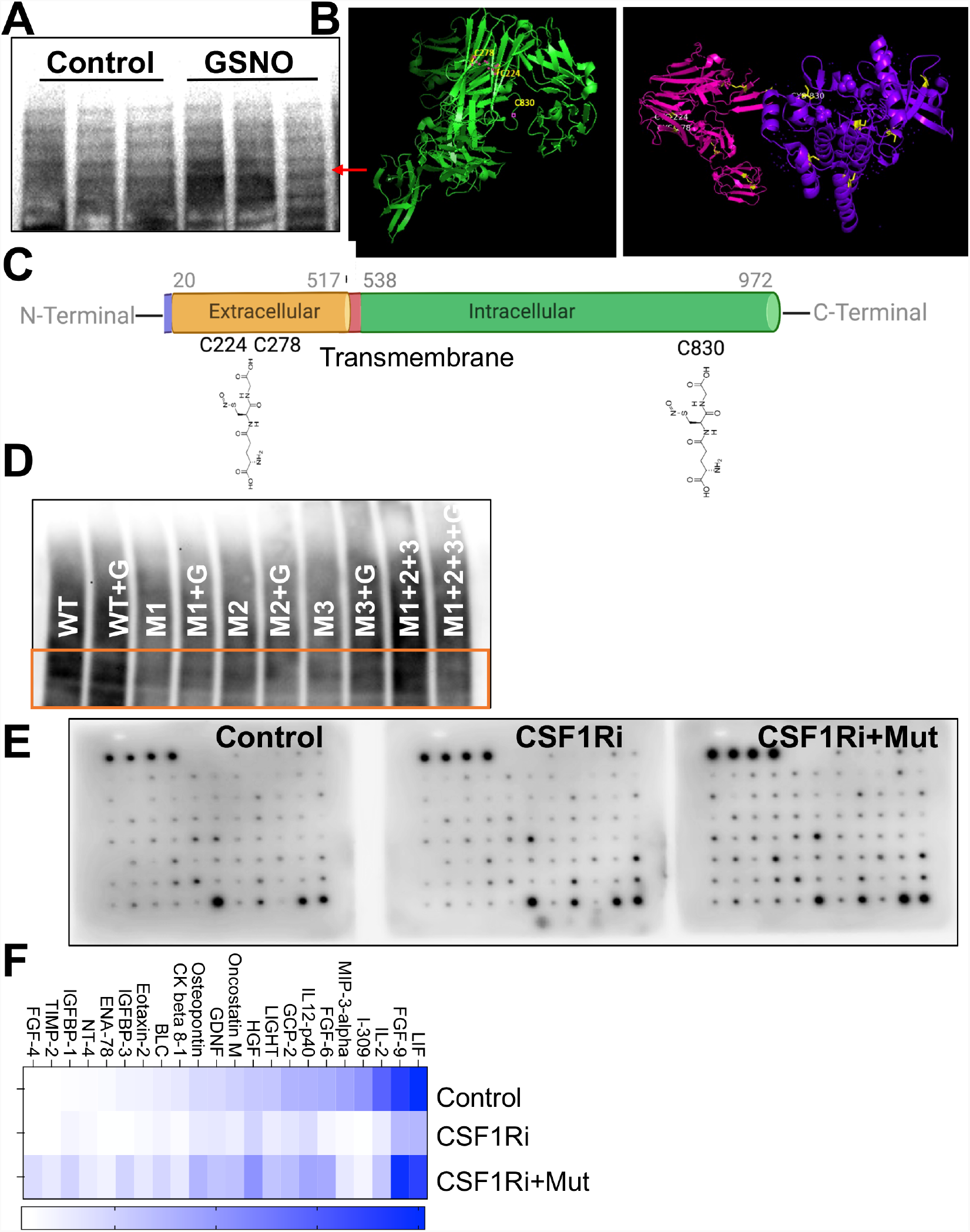
Increased Nitric oxide induces S-nitrosylation of CSF1R and augment the action of CSF1R inhibition to suppress PCa in vitro. (A) S-nitrosylation in tumor lysates in mice xenografts treated with vehicle (Control) and S-nitrosoglutathione (GSNO) for n=3 samples. (B) A 3D structure of CSF1R showing 3 cysteine sites highlighted in yellow with the highest threshold for nitrosylation. (C) A topology map of CSF1R showing intracellular and extracellular domains and the location of cysteine sites. (D) S-nitrosylation in cell lysates in 22Rv1 cell transfected with wild type, mutants with deletion at sites C224, C278 and C830 in the presence/absence of 50um S-nitrosoglutathione (GSNO). (E) Cytokine array image of the 22Rv1 cell lysates under different experimental conditions-untreated control, cells treated with CSF1R inhibitor at 0.5um dose and cells transfected with all 3 mutants along with the CSF1Ri. (F) Heat map showing different cytokines differentially expressed in cell lysates treated with CSF1Ri and CSF1R mutants with CSF1Ri. Data is presented as mean ± SEM. ***P < 0.001; **P < 0.01.

Next, to confirm if S-nitrosylation of CSF1R at Cys224, Cys278, and Cys830 is relevant for effective reduction of tumor promoting cytokines, we transfected 22RV1 cells with CSF1R plasmids (wild type and three mutants) and exposed the cells to vehicle or CSF1R blockade (GW2580, 0.5 μM) for 48 hours. Post treatment, protein lysates were subjected to cytokine antibody array. Results (Figure 6E) showed that mutating S-nitrosylation sites on CSF1R prevented effective inhibition of several pro-tumorigenic markers upon CSF1R inhibition such as LIF, FGF-9, IL-2, FGF-6, Oncostatin, GDNF, BLC, IFGBP-3 etc. (Figure 6F). Further, to study if S-nitrosylation of CSF1R is important for GSNO-induced changes on macrophage polarization, we transfected 22RV1 cells with the CSF1R plasmids and exposed the cells to vehicle or GSNO for 48 hours. 22RV1 cells transfected with wild type CSF1R plasmid only and treated with GSNO demonstrated reduced expression of androgen receptor (AR)(Suppl. Fig. 6B)^36^. Together, these results, along with our previous findings ^12, 39, 40, 41^ articulate the potential role of GSNO as a combinatorial partner to augment the anti-tumor effects of CSF1R inhibition against CRPC.

### Increased NO levels augment the action of CSF1R inhibition in suppression of PCa

Next, we tested the impact of combining CSF1Ri with GSNO on LNCAP and 22RV1 cells. Results showed, increasing concentrations of CSF1Ri combined with 50uM of GSNO concentration, significantly decreased cell proliferation in LNCAP cells (Supplementary Figure 7A). In 22RV1 cells, GSNO had minimal effects on cell proliferation ^12^(Supplementary Figure 7B) but increased the inhibitory impact of CSF1Ri on the expression of TMRPSS2 and PSA at RNA (Supplementary Figure 7C) and AR at protein levels (Supplementary Figure 7D).

Next, we used U937 cells and treated them with PMA in the presence of pro or anti-inflammatory macrophage inducing cocktails (Figure 7A). These cells were then exposed to GSNO (50uM) and CSF1Ri (0.5 μM) combination (NO-CSF1Ri). Results showed a significant increase in the population of M1 (CD68^+^) and a reduction in M2 (CD206^+^CD163^+^) macrophages, 48 hours post treatment (Figure 7A) suggesting that NO-CSF1Ri combination could effectively inhibit macrophage polarization in-vitro.

**Fig 7:**
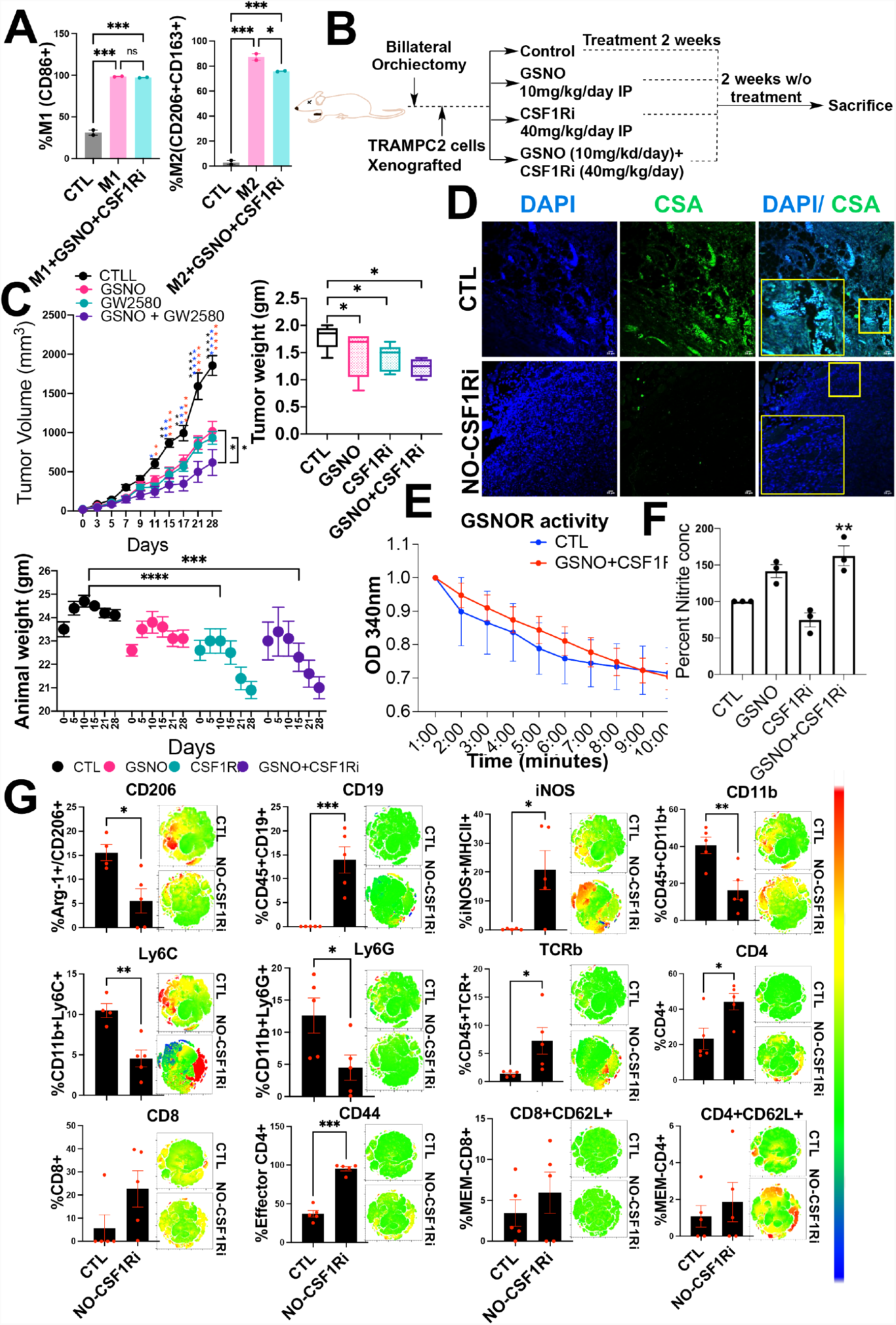
Impact of NO-CSF1Ri therapy in immune competent murine models. (A) Percentage of M1 (CD86+) and M2 (CD206+CD163+) macrophages in U937 cells used as a model for macrophage differentiation using specific M1 and M2 cocktails when treated in the presence/absence of GSNO+CSF1Ri combination. Data is represented as mean ± SEM. ***P < 0.001; **P < 0.01, *P < 0.05. ns=not significant. (B) Experimental plan. (C) Tumor volume, animal weight, and tumor weight represented for n=5 animals per treatment condition. (D) Representative immunofluorescent images for tumor sections for mice treated with vehicle and a combination of GSNO and CSF1R inhibitor (NO-CSF1Ri) and stained with 4′,6-diamidino-2-phenylindole (DAPI) (blue) and Cysteine sulfinic acid (red) (n=3). (E) GSNOR activity measured at 340nm for a span of 10 minutes using tumor lysates treated with vehicle and a combination of GSNO and CSF1R inhibitor (n=3). (F) Percent nitrite concentration under different treatment conditions as estimated using the Griess test in tumor lysates (n=3). (G) Immune-phenotyping done in tumor cells in untreated and NO-CSF1Ri mice (n=5) for various markers: CD206, CD19, iNOS, CD11b, Ly6C, Ly6G, TCRB, CD4, CD8, CD44 and CD62L respectively. Data are Mean ± SEM. ***P < 0.001; **P < 0.01, *P < 0.05.

Next, we examined the impact of NO-CSF1Ri combination on reducing tumor burden of CRPC murine models (castrated C57BL/6 mice, grafted with TRAMP-C2 cells). Treatment conditions included GSNO (10mg/kg/day IP), CSF1Ri (GW2580, 40mg/kg/day IP), or combination (GSNO at 10mg/kg/day + GW2580 at 40mg/kg/day IP) (Figure 7B). Importantly, the greatest reduction in tumor burden was achieved in mice receiving the NO-CSF1Ri combination (Figure 7C) compared to CSF1Ri or GSNO monotherapies. Animals’ weight was reduced in the combination and CSF1Ri monotherapy group when compared to the control group, unlike mice receiving GSNO alone (Figure 7C). The tumor grafts from combination group showed reduced oxidation pockets (Figure 7D). Moreover, GSNOR activity was reduced by NO-CSF1Ri treatment (Figure 7E). Additionally, Griess test on tumor grafts showed that nitrate levels were significantly high (**p≤0.01) in tumors from NO-CSF1Ri treatment group (Figure 7F). This suggests, offsite (intraperitoneal), treatment with GSNO as a monotherapy or in combination with CSF1Ri is capable to increase the nitrite ions in the tumor grafts.

Moreover, NO-CSF1Ri treated mice exhibited reduced expression of AR, ARV7, PSA, TMRPSS2, p-GSK, p-ERK, CD206, KI67 and p90RSK (Supplementary Figure 7E-G). Furthermore, cytokine array showed, NO-CSF1Ri combination reduced pro-tumorigenic cytokines such as ENA-78, BLC, IP-10, Oncostatin M, FGF-6, IGFBP-3, RANTES, Osteopontin, GCP-2, IL-15, MCP-4, NT-4, TGF-b 3, IL-3, SCF, IL12-p40, IGFBP-4, FGF-4, SDF-1, PDGF-BB, IL-1beta, PARC, TPO, IFN-gamma, FGF-7, and IL-2 (Supplementary Figure 8A). Next, Immunophenotyping demonstrated that NO-CSF1Ri combination significantly decreased intratumoral percentage of M2 macrophages (CD45^+^CD206^+^), myeloid cells (CD45^+^CD11b^+^), MDSCs (CD45^+^LY6C^+^), and PMN-MDSCs (CD45^+^LY6G^+^) and increased the percentage of M1 macrophages (iNOS^+^MHCII^+^), cytotoxic T lymphocytes (CD8^+^), and effector T cells (CD44^+^CD62L2^-^) (Figure 7G). Additionally, immunostaining using antibodies against iNOS (M1 macrophage marker), F4/80 and pERK ^42^, showed that the NO-CSF1Ri combination suppress the expression of F4/80 and pERK while inducing the expression of iNOS (Supplementary Figure 8B-C) to reduce macrophage polarization in vivo. Together, these results validate that exogenous supplementation of NO levels augment the action of CSF1Ri to suppress PCa.

## Discussion

We observed that eNOS uncoupling in high grade human PCa (CRPC) results in reduced NO levels and increased overall oxidation. The reduced NO levels negatively influences the anti-tumor effectiveness of CSF1R blockade therapy as CSF1R at Cys224, Cys278, and Cys830 sites could not be effectively S-nitrosylated. These observations strongly suggested the relevance of exogenous NO treatment on reverting the side effects of eNOS uncoupling and augmenting CSF1R blockade against CRPC. In our previous studies, we showed that increased levels of NO are associated with lowering luteinizing hormone and testosterone^41^, suggesting the tumor inhibitory role of NO against PCa^12, 43^. Here, our findings answered that- (a) uncoupled eNOS increases the overall oxidative stress in high grade PCa; (b) lack of S-nitrosylation of three cystine residues the ant-tumor abilities of CSF1R blockade; (c) exogenous NO treatment S-nitrosylates CSF1R at Cys224, Cys278, and Cys830 sites; (d) in the presence of exogenous NO, CSF1Ri blockade can effectively reduce CRPC tumor burden, compared to NO or CSF1R blockade monotherapies; (e) intratumoral percentage of M2 macrophages (CD45^+^F4/80^+^, CD206^+^), CD45^+^LY6G^+^, and CD45^+^LY6C^+^, CD45^+^CD11b^+^ cells are decreased and that of M1 macrophages (iNOS^+^MHCII^+^), cytotoxic T lymphocytes (CD8^+^), effector T cells (CD44^+^CD62L2^-^), B cells (CD45^+^CD19^+^) are increased upon NO-CSF1R blockade; (e) several pro-tumorigenic cytokines which are not effectively reduced by CSF1Ri monotherapy are reduced upon NO-CSF1R blockade. Together the results suggested that exogenous increase in NO could increase the efficacy of CSF1R blockade against CRPC

Our study has limitations that should be addressed with further work. For example, 1. though we tested different combinations of GSNO and CSF1Ri *in vitro*, the correlation between concentrations of NO-CSF1Ri and in-vivo tumor burden reduction remains unexplored. 2. We showed that S-nitrosylation of three different cystines on CSF1R molecule via GSNO is important to modulate inhibitory effects on CRPC tumors, however, what other molecules are S-nitrosylated and how they could modulate NO-CSF1R blockade therapy still needs to be explored. We showed earlier that increased NO levels are capable of affecting hypogonadism ^41^ and leading to CRPC tumor suppression, which therefore leads to the possibility that tumor inhibition could be profound in non-castrate mice. However, this assumption still needs to be explored.

To conclude, our findings present insights into an underexplored area of CRPC therapeutics and the mechanistic insights gathered in this study provides a strong rational for initiating a clinical trial with NO-CSF1R blockade in patients with advanced metastatic stages of cancer.

## Acknowledgments

We thank all the mentors (Dr. Dipen J Parekh), collaborators (Dr. Bonnie Bloomberg), graduate student (Anastasia Vedenko) and Postdoctoral students (Dr. Deepa Seetharam) for their insights, suggestions, and support during this study.

## Availability of Data and Materials

All data, code, and materials used in the study is available to any researcher for purposes of reproducing or extending the analysis. Materials transfer agreements (MTAs) will be required to request the access. RNA sequencing data has been deposited in a public database.

