## Supplementary material for "Nitric oxide S-nitrosylates CSF1R to augment the action of CSF1R inhibition against castration resistant prostate cancer": Supp Material

**This PDF file includes:**

Materials and Methods
Supplementary Text
Figs. S1 to S8
Tables S1 to S3
Raw Blots

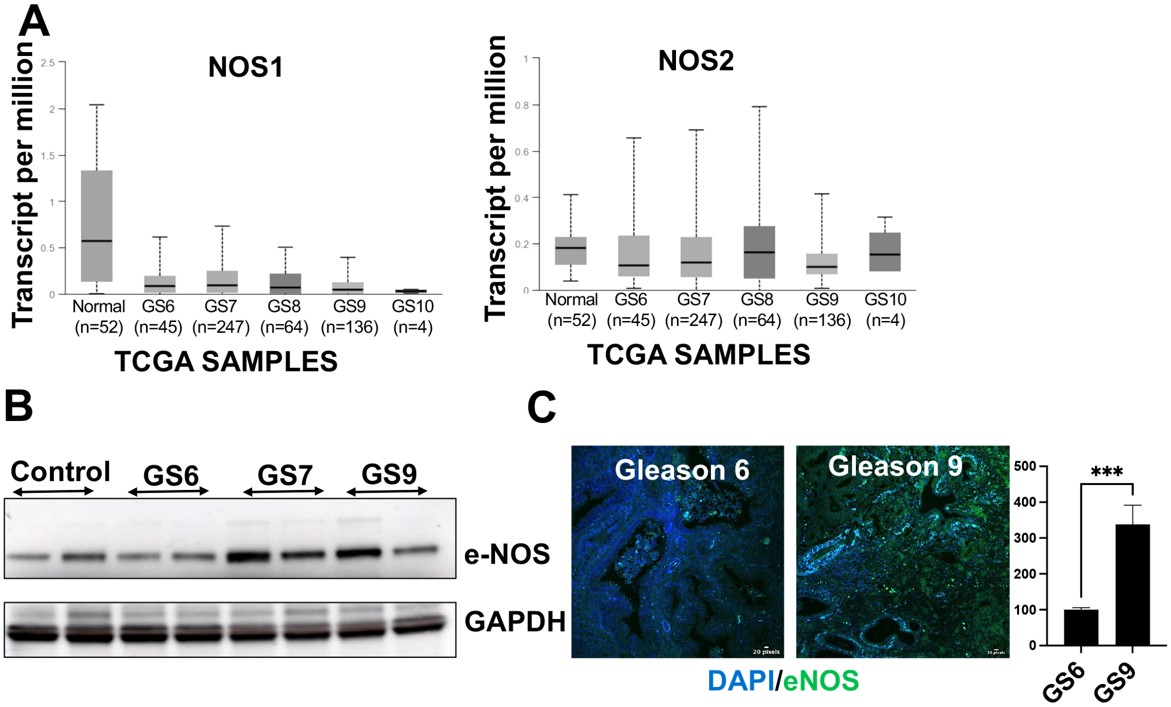


**Supp Fig 1:** (A) Graphs showing the relative expression of NOS1 and NOS2 across different Gleason Grades as studied using RNA sequencing data for Prostate adenocarcinoma from The Cancer Genome Atlas (TCGA). (B) Western blot following immuno-precipitation of Peripheral Blood Mononuclear Cells (PBMCs) showing the expression of NOS3 (eNOS) (n=2). (C) Immunofluorescent staining images showing expression of eNOS (green) in patient biopsies form Gleason Grade 6 and 9.

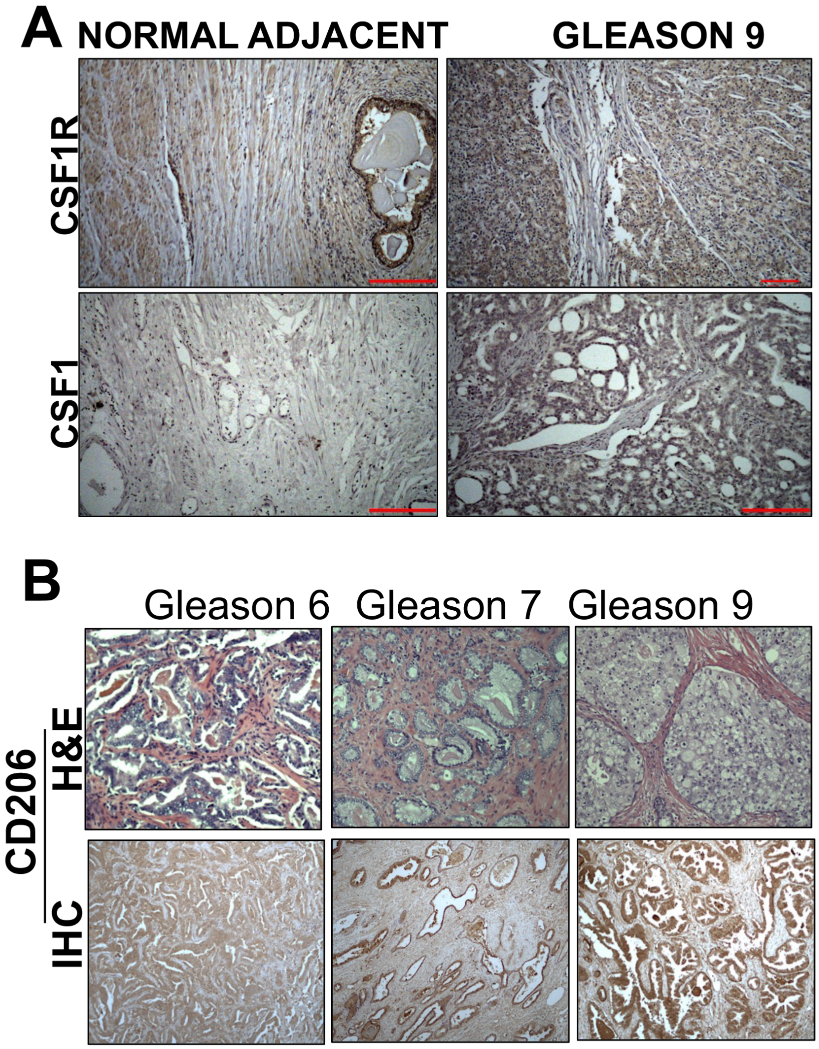


**Supp Fig 2:** (A) Representative images for Hematoxylin and DAB staining showing expression of CSF1R and CSF1 in Normal adjacent vs Gleason grade 9 patient biopsies. Graphs shiwing percent DAB intensity for relative CSF1 and CSF1R expression. (B) Representative immunohistochemical as well as Hematoxylin and eosin staining images showing expression of CD206 (M2 macrophage marker) in patient biopsies from different Gleason Grades i.e., 6, 7 and 9.

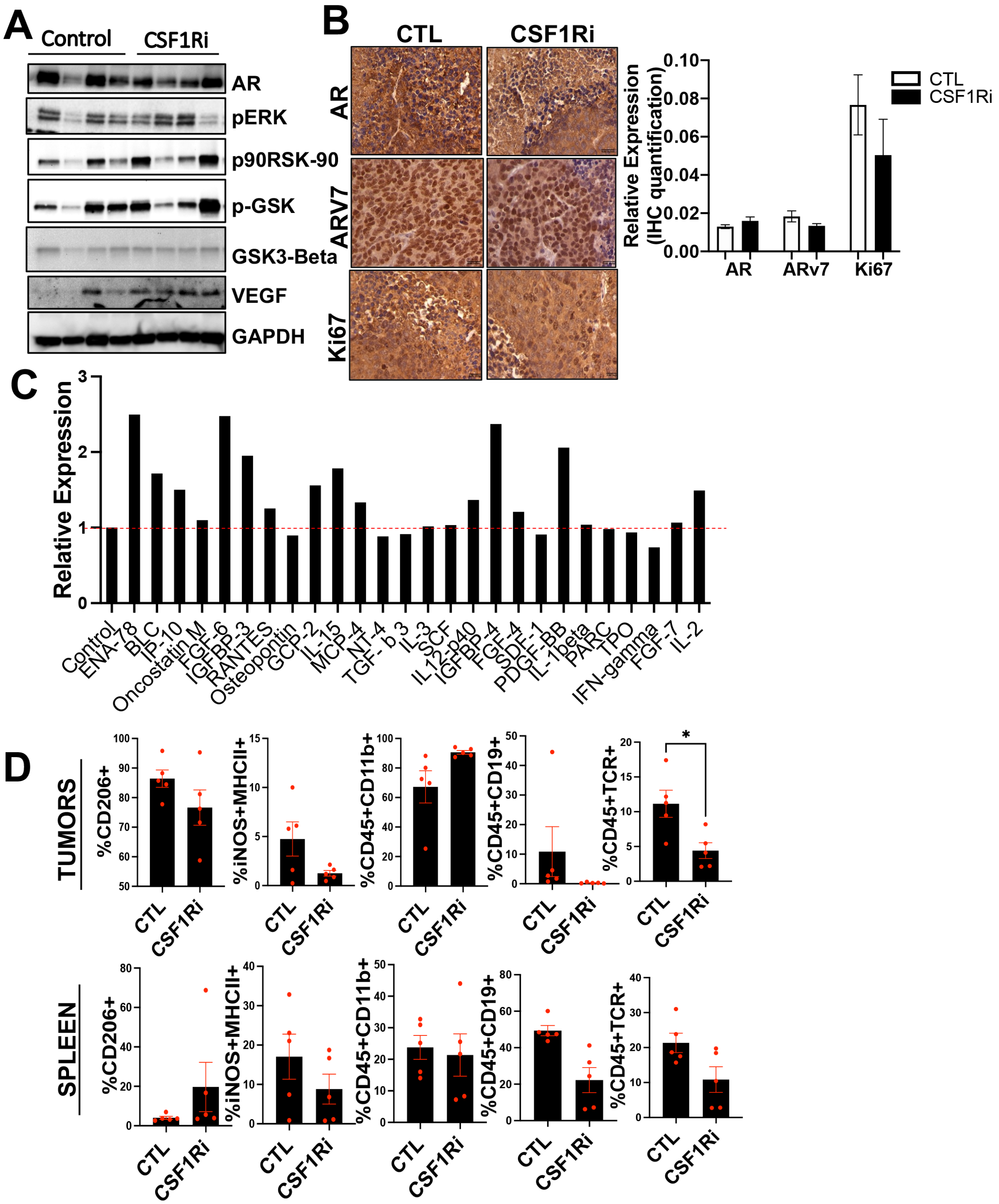


**Supp Fig 3: (A)** Protein levels of AR, pERK, p90-RSK, pGSK, GSK3 BETA, VEGF and GAPDH in tumor grafts treated with CSF1R inhibitor (GW2580). (B) Representative IHC images with the quantification showing expression of AR, ARv7 and KI67 using DAB staining in tumor sections. (C) Cytokine antibody array showing selective tumor promoting candidates whose relative expression was induced or minimally changed upon CSF1R inhibition in tumor grafts. (D) Graphs showing differential immune population markers in spleenocytes.

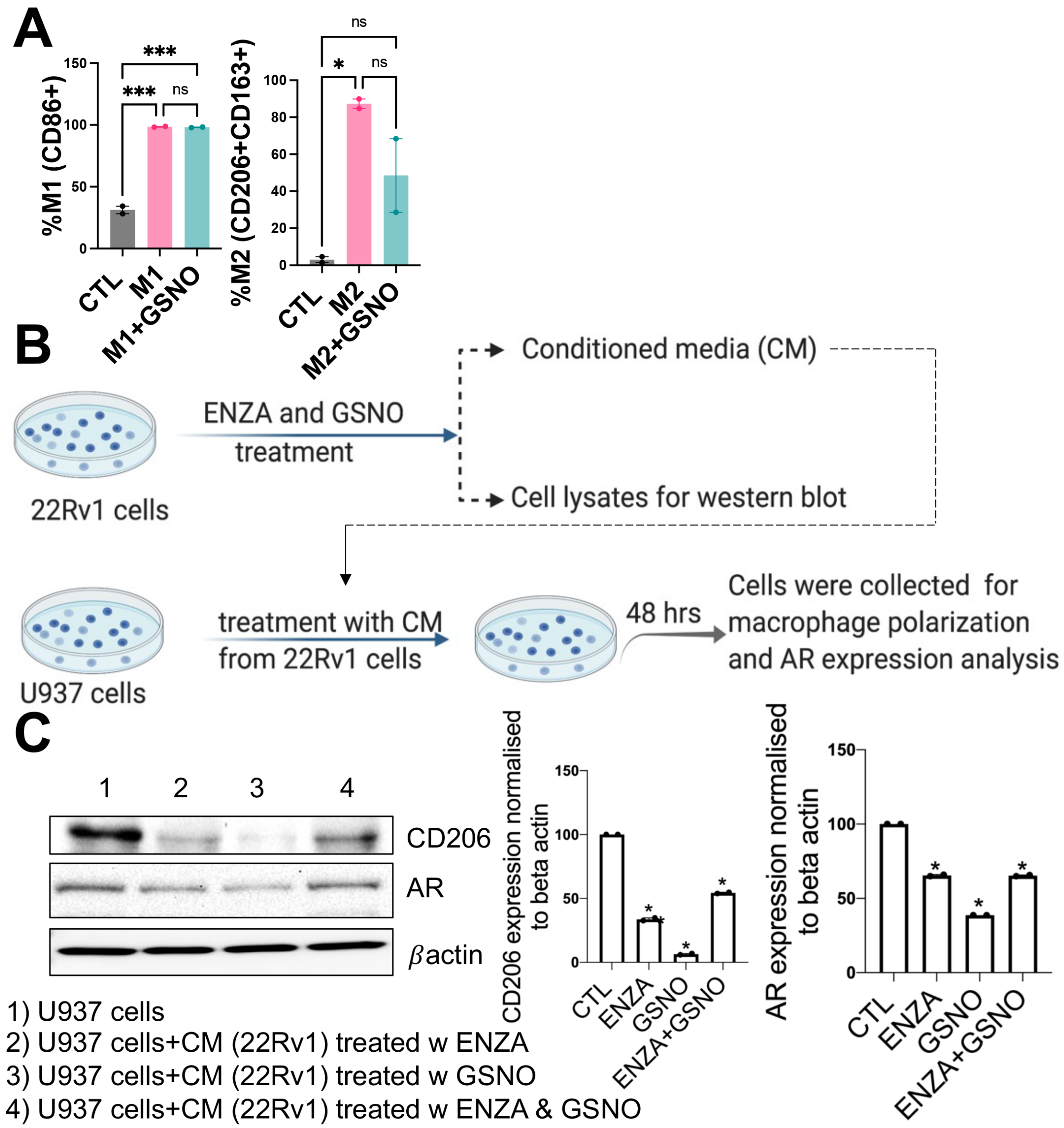


**Supp Fig 4:** (A) Percentage of M1 (CD86+) and M2 (CD206+CD163+) macrophages in U937 cells used as a model for macrophage differentiation using specific M1 and M2 cocktails when these cells were treated in the presence/absence of GSNO. (B) Experimental plan showing the course of treatment in U937 cells when these are exposed to conditioned media collected from 22Rv1 cells exposed to AR antagonist (Enzalutamide) with/without GSNO. (C) Western blot analysis and quantification for AR and CD206 expression done in U937 cells treated with different treatment conditions.

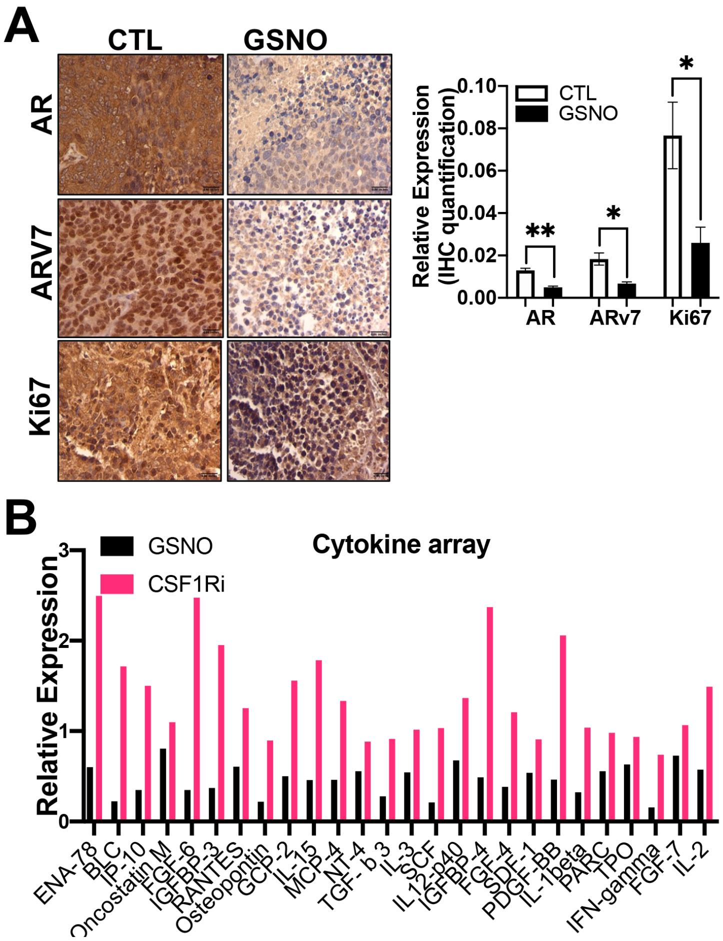


**Supp Fig 5:** (A) Representative IHC images as well as quants for AR, ARv7 and Ki67 expression as estimated in tumor sections treated with vehicle and GSNO. (B) Cytokine antibody array showing selective tumor promoting candidates whose relative expression was reduced or minimally changed upon GSNO treatment as compared to CSF1R inhibitor treated tumor grafts.

A

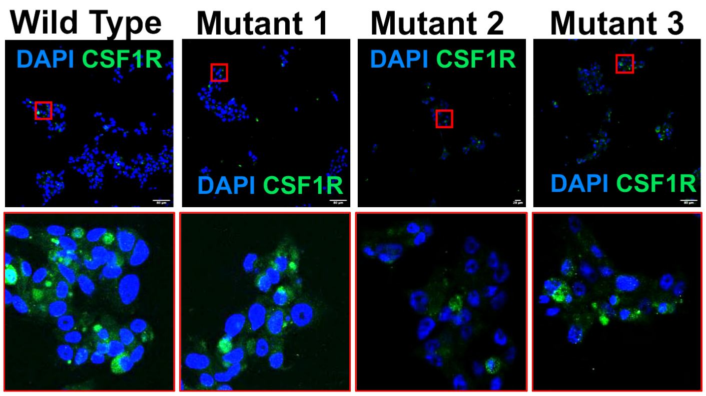


B

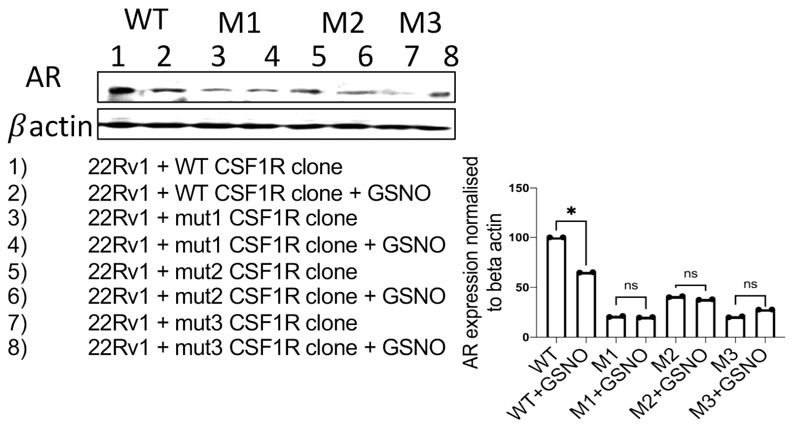


**Supp Fig 6:** (A) Representative images showing sub-cellular localization of wild type and three CSF1R mutants (M1, M2 and M3) with deletions at C224, C278 and C830 cysteine residues in 22Rv1 cells. (B) Analysis of AR expression in 22Rv1 cells transfected with WT and 3 mutants in the presence and absence of GSNO.

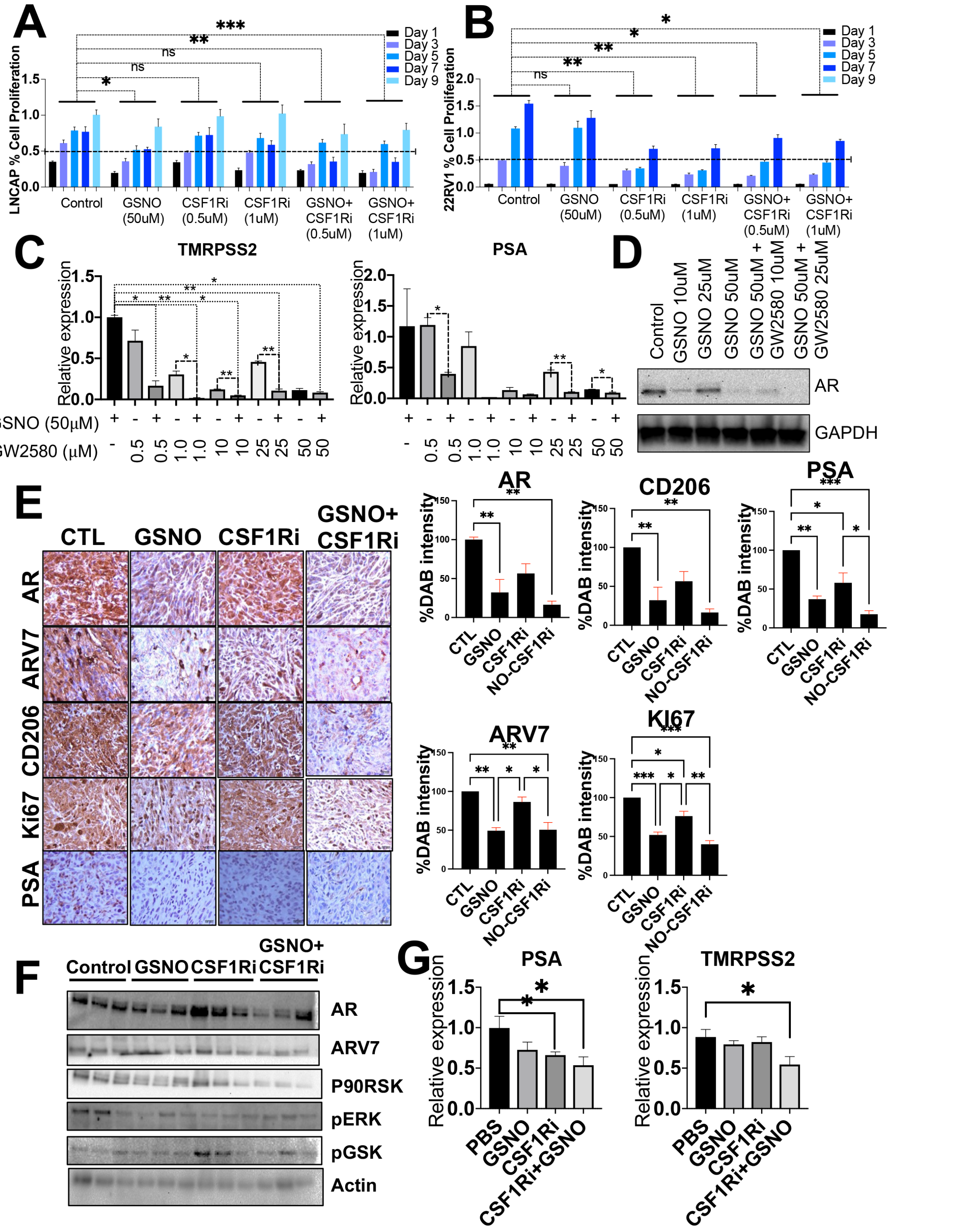


**Supp Fig 7:** (A) Impact of increasing concentrations of GW2580 and fixed concentration of GSNO (50uM) evaluated on LNCAP and (B) 22RV1 cell proliferation using MTT assay (C) RNA levels of

TMRPSS2 and PSA under different treatment conditions. (D) Protein levels of AR in 22Rv1 cells treated with variable doses of GSNO and CSF1R inhibitor (GW2580). (E) Western blot analysis for checking the levels of AR, ARv7, p90RSK, pERK, and pGSK in tumor grafts treated with vehicle, GSNO (10 mg/kg), GW2580 (40 mg/kg) and a combination of both GSNO and GW2580 respectively (n=3). Beta actin was used as a loading control. (F) mRNA levels of PSA and TMPRSS2 estimated in tumor grafts. Data is presented as mean ± SEM.

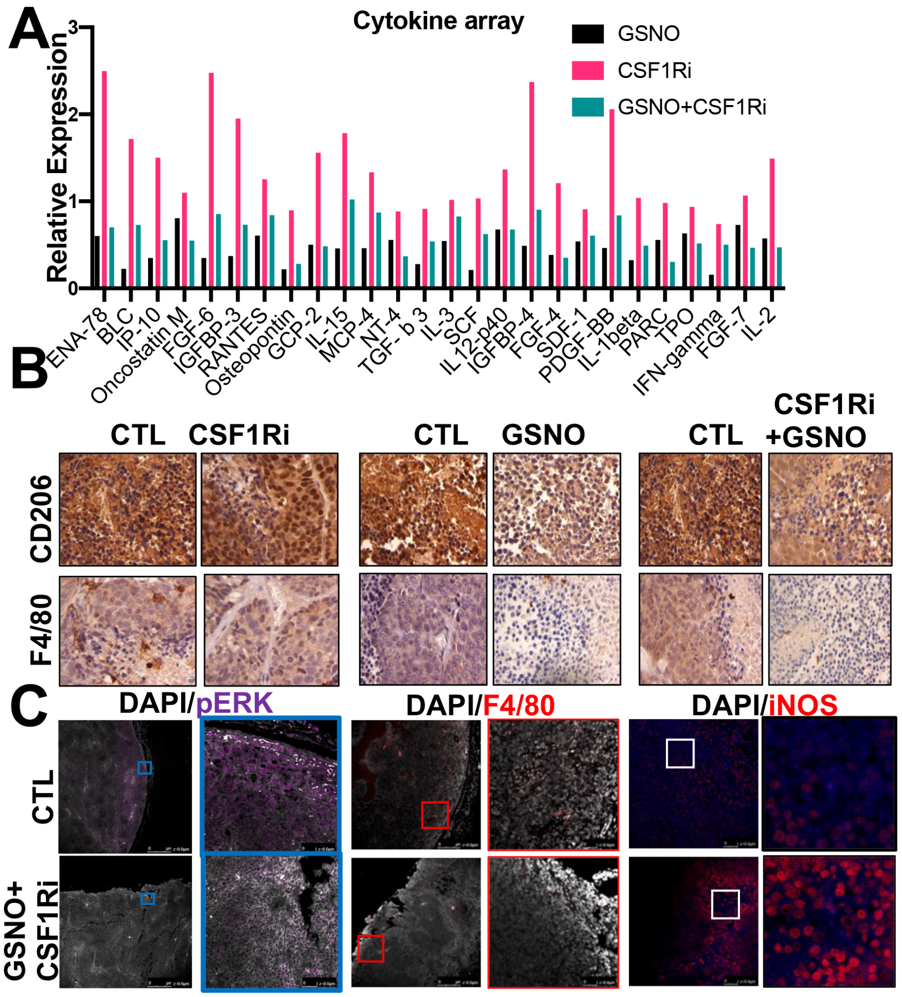


**Supp Fig 8:** (A) ) Cytokine antibody array showing selective tumor promoting candidates whose relative expression was reduced/induced upon GSNO, CSF1R as well as combination treatment as estimated in 22Rv1 tumor grafts. (B) Immunohistochemistry images showing relative expression using DAB staining for the expression of AR, ARv7, CD206, KI-67, and PSA in TRAMPC2 mice exposed to GSNO, CSF1Ri, and NO-CSF1Ri treatments. (C) Immunohistochemical images showing the expression of M2 macrophage markers-CD206 and F4/80 in 22Rv1 tumors (D) Representative immunofluorescence images showing the impact of NO-CSF1Ri treatment on the expression of pERK, F4/80, and iNOS in 22Rv1 xenografts.

| Cytokine | Associated Cancers | Impact on Tumor Microenvironment | PMID |
|---|---|---|---|
| *ENA-78* | Intrahepatic cholangiocarcinoma<br>Hepatocellular carcinoma<br>renal cell carcinoma | promoting tumor progression by regulating the functions of various host cells | 24293410 |
| *BLC* | Lymphoma<br>gastric lymphomas | Involvement in Tertiary Lymphoid Structure Formation in Tumors | 19955043<br>31354634 |
| *IP-10* | pancreatic cancer | promote immunosuppression and tumor progression | 26405599 |
| *Oncostatin M* | Pancreatic Ductal Adenocarcinoma<br>Skin squamous cell carcinoma | promote epithelial–mesenchymal plasticity, and acquisition of cancer stem cell (CSC) properties | 28288136<br>30559930 |
| *FGF-6* | Prostate cancer | Promote motility and invasiveness of a variety of cancers | 19508171 |
| *IGFBP-3* | esophageal squamous cell carcinoma | Promotes epithelial-mesenchymal transition | 24482736 |
| *RANTES* | breast cancer | promotes cancer growth by suppressing oxidative stress in hypoxic tumor microenvironment | 32478064 |
| | Hodgkin Lymphoma<br>Prostate Cancer<br>Breast cancer | promoted the migration and invasion | 24523569<br>31578575<br>31578575 |
| *Osteopontin* | Hepatic cancer | activate TAMs and influence angiogenesis | 24657887 |
| *GCP-2* | Prostate Cancer | regulation of tumor angiogenesis | 30591657 |
| | Breast cancer | serve as a proto-oncogene in the tumor microenvironment | 27384993 |
| *IL-15* | Breast cancer | promoting antitumor responses | 30587590<br>30587590 |
| *MCP-4* | lung carcinoma<br>Prostate cancer | promotes recruitment of monocytes and macrophages to tumor sites | 30483271<br>16705739 |
| *NT-4* | Brain cancer | Promotes endogenous neuron growth | 31900168 |
| *TGF-β* | Hepatocellular carcinoma | induces malignant MECs to undergo EMT | 28624577 |
| | Breast cancer | potent inducer of tumor angiogenesis | 28624577 |
| *SCF* | prostate cancer | Mast cell infiltration and activation in tumors | 18524989<br>28656507 |
| *IL12-p40* | prostate cancer | generation of T-helper type 1 cells | 29073075 |
| *IGFBP-4* | Pancreatic Cancer<br>Lung cancer | inhibits cell growth, migration and invasion | 32414222<br>28150906 |
| *FGF-4* | Ovarian cancer | promotes tumor initiation through autocrine and paracrine mechanisms | 25329002 |
| *SDF-1* | Multiple myeloma | angiogenesis, osteoclastogenesis or tumor cell migration and adhesion to stromal cells | 26655999 |
| *PDGF-BB* | Colorectal and pancreatic cancer | Promotes epithelial to mesenchymal transition | 17641778 |
| *IL-1beta* | Breast cancer | Promotes immunosuppression | 30545915 |
| *PARC* | Breast cancer | recruitment of naive T-cells and dendritic cells | 26416449 |
| *TPO* | Thyroid cancer | Facilitates Progression | 27942419 |
| *IFN-gamma* | Breast cancer<br>Fallopian cancer | Hypoxia regulation in tumor microenvironment | 20457620<br>20974954 |
| *FGF-7* | gastric cancer | promotes invasion and migration | 28339036 |
| *IL-2* | Cervical Cancer | critical for normal lymphocyte proliferation | 27293315 |

**Supplementary Table 1**: Table showing list of cytokines involved in various cancers and their role in modulating tumor microenvironment.

| Target | Fluoro | Company | Catalog# |
|---|---|---|---|
| Viability | Live/dead Blue | Thermofisher | Cat # L34962 |
| Ly6G | PE-Cy5 | ThermoFisher | Cat # 15-9668-82<br>Lot# 2283652 |
| CD19 | PE-Cy5.5 | ThermoFisher | Cat # 35-0193-82<br>Lot# 2317034 |
| iNOS | PE-Cy7 | ThermoFisher | Cat # 25-5920-82<br>Lot# 2228654 |
| Arg1 | APC | ThermoFisher | Cat # 17-3697-82<br>Lot# 2079508 |
| MHCII | APC-Fire 750 | ThermoFisher | Cat # 58-5321-82<br>Lot# 2338622 |
| FOXP3 | PE | ThermoFisher | Cat # 12-5773-82<br>Lot# 2344844 |
| Ki67 | APC-eF780 | ThermoFisher | Cat # 47-5698-82<br>Lot# 2311269 |
| Ly6C / Ly6G | BUV563 | BD | Cat # 741226<br>Lot# 1132321 |
| TCRb | BUV496 | BD | Cat # 749914<br>Lot# 1132322 |
| CD45 | BUV805 | BD | Cat # 741957<br>Lot# 1132327 |
| CD62L | BUV563 | BD | Cat # 741230<br>Lot# 1132332 |
| CD44 | BUV737 | BD | Cat # 612799<br>Lot# 295805 |
| PD-L1 | BV421 | Biolegend | Cat # 124315<br>Lot# B278497 |
| CD206 | BV650 | Biolegend | Cat # 141723<br>Lot# B313883 |
| CD11b | BV750 | Biolegend | Cat # 101267<br>Lot# B327959 |
| F4 / 80 | AF488 | Biolegend | Cat # 123120<br>Lot# B272102 |
| CCR7 | PE | Biolegend | Cat # 120106 |
| MHCII | APC-Fire 750 | Biolegend | Cat # 107652 |

**Supplementary Table 2**. Shows antibodies that were used in immunophenotypic panel to study the impact of treatments on several immune cells.

For >sp|P07333.2|CSF1R_HUMAN
GPS-SNO Data

| Position (ALL) | Peptide | Score | Cutoff | Cluster |
|---|---|---|---|---|
| 42 | GATVTLRCVGNGSVE | 2.19 | 0 | Cluster B |
| 84 | QNTGTYRCTEPGDPL | 1.223 | 0 | Cluster B |
| 127 | DQDALLPCLLTDPVL | 18.715 | 0 | Cluster C |
| 177 | IQSQDYQCSALMGGR | 1.228 | 0 | Cluster B |
| 224 | GEAAQIVCSASSVDV | 3.304 | 0 | Cluster B |
| 278 | QHAGNYSCVASNVQG | 2.473 | 0 | Cluster B |
| 419 | NGSGTLLCAASGYPQ | 1.141 | 0 | Cluster B |
| 434 | PNVTWLQCSGHTDRC | 0.87 | 0 | Cluster B |
| 441 | CSGHTDRCDEAQVLQ | 0.148 | 0 | Cluster A |
| 485 | EHNQTYECRAHNSVG | 0.459 | 0 | Cluster A |
| 522 | FTPVVVACMSIMALL | 0.683 | 0 | Cluster A |
| 653 | IVNLLGACTHGGPVL | 1.197 | 0 | Cluster A |
| 666 | VLVITEYCCYGDLLN | 0.727 | 0 | Cluster A |
| 667 | LVITEYCCYGDLLNF | 1.951 | 0 | Cluster B |
| 774 | AFLASKNCIHRDVAA | 2.402 | 0 | Cluster B |
| 830 | APESIFDCVYTVQSD | 2.478 | 0 | Cluster B |
| 892 | IYSIMQACWALEPTH | 1.459 | 0 | Cluster A |
| 907 | RPTFQQICSFLQEQA | 0.12 | 0 | Cluster A |
| 953 | SSSEHLTCCEQGDIA | 1.94 | 0 | Cluster B |
| 954 | SSEHLTCCEQGDIAQ | 0.918 | 0 | Cluster A |
| 972 | QPNNYQFC******* | 17.277 | 0 | Cluster C |

| Position High Thresl | Peptide | Score | Cutoff | Cluster |
|---|---|---|---|---|
| 224 | GEAAQIVCSASSVDV | 3.304 | 2.443 | Cluster B |
| 278 | QHAGNYSCVASNVQG | 2.473 | 2.443 | Cluster B |
| 830 | APESIFDCVYTVQSD | 2.478 | 2.443 | Cluster B |

**Supplementary Table 3**. Shows potential cystine sites on CSF1R which could be S-nitrosylated by NO. A total of 20 cystine sites were identified and the information of predicted threshold for being S-nitrosylates shown.
